# Charting the transcriptomic landscape of primary and metastatic cancers in relation to their origin and target normal tissues

**DOI:** 10.1101/2023.10.30.564810

**Authors:** Neel Sanghvi, Camilo Calvo-Alcañiz, Padma S. Rajagopal, Stefano Scalera, Valeria Canu, Sanju Sinha, Fiorella Schischlik, Kun Wang, Sanna Madan, Eldad Shulman, Antonios Papanicolau-Sengos, Giovanni Blandino, Eytan Ruppin, Nishanth Ulhas Nair

## Abstract

Metastasis is a leading cause of cancer-related deaths, yet understanding how metastatic tumors adapt from their origin to target tissues is challenging. To address this, we assessed whether primary and metastatic tumors resemble their tissue of origin or target tissue in terms of gene expression. We analyzed gene expression profiles from various cancer types, including single-cell and bulk RNA-seq data, in both paired and unpaired primary and metastatic patient cohorts. We quantified the transcriptomic distances between tumor samples and their normal tissues, revealing that primary tumors are more similar to their tissue of origin, while metastases shift towards the target tissue. Pathway-level analysis highlighted critical transcriptomic changes during metastasis. Notably, primary cancers exhibited higher activity in cancer hallmarks, including *Activating Invasion and Metastasis*, compared to metastatic cancers. This comprehensive landscape analysis provides insight into how cancer tumors adapt to their metastatic environments, providing a transcriptome-wide view of the processes involved.

## INTRODUCTION

Most cancer deaths are caused by metastasis, a hallmark of cancer that remains poorly understood. The metastatic process, in which primary tumor cells adapt and travel from their origin tissue to target tissue(s), is a complex cascade of molecular alterations^1^. Numerous studies that have examined the molecular features of metastatic cancers, highlighting hallmarks such as cell motility and invasion, the ability to modulate the target tissue, phenotype plasticity, and the ability to colonize the target tissue^2^. In particular, metabolic adaptations play an important role in metastases as they seed in their target tissue^3^, including altered fatty acid metabolism and changes in energy supply to initiate and promote metastasis^4–7^. Primary tumors, on the other hand, have been found to remain overall metabolically similar in their expression profiles to their normal origin tissues, though with some upregulation of pathways such as nucleotide synthesis and glycolysis^8^, suggesting different genomic rewiring during growth from metastases.

Previous work has directly compared primary and metastatic tumors’ transcriptomics, identifying differentially expressed markers and metastatic alterations across various cancer types^3,9,10^. Furthermore, genomic sequencing of 25,000 primary and metastatic tumors revealed higher chromosome instability, fractions of clonal mutations, and enrichments in drug resistance mechanisms in metastases compared to primary tumors^11^. However, genomic or transcriptomic rewiring during metastasis is also dependent on the target tissues in the metastatic process. Metastatic tumors are known for their particular organotropism, i.e., their specific spread to distant metastatic sites, with different cancer types displaying distinct distributions of target tissues of metastasis^10^. On this notion, Ngyuen & Fong et al^11^ linked specific genomic alterations to the particular organotropisms in various cancer types, suggesting the importance of the target tissue in the metastatic process. However, the origin tissues too play in role, as differential expression analysis of primary tumors from The Cancer Genome Atlas Network^12^ (TCGA) and adjacent normal tissues exhibited heterogeneity of tricarboxylic acid cycle, anaplerotic reactions, and electron transport chain expression profiles, with unique expression changes specific to tumors from different origin tissues^13^. Thus, both the origin and target tissues of a tumor influence growth and adaptations during metastasis.

There has also been considerable work on this interplay of tumors and their corresponding normal tissues. Lee et al^14^ sequenced five sets of paired normal and primary/metastatic samples of colorectal cancer patients with liver metastases, finding cell cycle, mitosis, and cell division pathways to be significantly enriched in both the primary carcinoma and metastases, with the tumors having similar expression overall. Comparison of single-cell transcriptome data of colorectal tumors (paired primary and metastatic) and paired normal origin and target tissues found PPAR signaling pathway-related genes upregulated in tumor epithelial cells compared to normal ones. Inhibiting this signaling pathway resulted in stunted colorectal tumor organoid growth *in vitro*, suggesting it as a tumor-specific target for colorectal cancer^15^. Metastasis to specific target sites from various cancer types has also been studied through single-cell atlases, with the brain and liver being common sites, finding organ-specific markers of metastasis in normal tissues and contributing to the burgeoning understanding of tumor organotropism^16–18^. Roshanzamir et al^19^ corroborate this notion, having utilized bulk transcriptomic data and metabolic network analyses of triple negative breast cancer (TNBC) metastases to find that the metastases adapt their metabolism to various target tissues while also retaining metabolic signatures associated with the primary TNBC. They found enrichment of metabolic pathways, e.g., bile acid metabolism, and immune response functions, e.g., coagulation, in metastatic tumors compared to primary tumors, while primary tumors were enriched more with proliferation, signaling, and inflammatory response pathways. Furthermore, their genome-scale metabolic model analysis of these TNBC metastases, their primary tumors, and healthy tissues highlighted the importance of several metabolic pathways like nicotinate and bile acid metabolism as potential pathways for targeting metastases.

With the growing interest in studying the molecular alterations occurring in cancer cells as they seed and adapt in their metastatic niche, we sought to identify *pan-cancer* patterns of transcriptomic similarities and differences to noncancerous origin and target tissues, which has not yet been characterized for primary and metastatic tumors in a *genome-wide* manner. Here we conduct such a study, performing a large-scale, pan-cancer gene expression comparison between (1) a tumor’s origin tissue, (2) the primary tumor, (3) the metastatic tumors, and (4) their target tissues. Studying this fundamental research question at this time is a non-trivial challenge as, to date, a large-scale, pan-cancer transcriptomic dataset containing per-patient fully paired sets of normal tissue, primary tumors, and metastatic tumors does not exist. Yet, we believe that it is timely to begin exploring this question as best as its current feasible with the currently available data. To this end, we set out to conduct a few analyses in concert. We begin by comparing the relevant entities in a small quadruple-paired cohort (all four sample types from same patient), then analyze multiple paired (primary and metastatic only) cohorts, and two large-scale unpaired cohorts of bulk transcriptomics, along with a paired single-cell transcriptomics dataset and computationally deconvolved bulk dataset. Based on these data, we aim to identify the underlying alterations that drive metastasis across cancer types in a robust, consistent manner as possible. A summary of all datasets used in this work is provided in **Table S1A**.

Given these data sets, we asked three basic research questions: (1) First, are primary and metastatic tumors transcriptionally more similar to the organ from which they originate (termed the *origin tissue*) or to the organ to which they metastasize (termed the *target tissue*)? (2) Second, on a pathway level, which pathways’ expression in metastatic tumors is most similar to their target tissue for a given primary cancer type? (3) Third, are primary cancers primed for metastasis before leaving the tissue of origin? This knowledge can provide a genome-wide view on how cancer cells adapt to their target organ microenvironment; pathways closer to the target tissues in metastatic tumors but not in primary ones point to a metastatic niche adaptation, while pathways closer to the origin tissues in metastatic tumors suggest conserved oncogenic transcriptomic signatures.

## RESULTS

### Overview of the Analysis

To study whether the overall gene expression of primary and metastatic tumors is closer to their tissue of origin or to their target tissue, we acquired datasets from publicly available sources such as the Gene Expression Omnibus (GEO), The Cancer Genome Atlas^12^ (TCGA), and the Genotype-Tissue Expression^20^ (GTEx) atlas, sourcing the gene expressions for four key entities: **(a)** *primary tumors* either annotated as having metastases or having paired metastases from the same patient, **(b)** *metastatic tumors*, **(c)** normal, non-cancerous tissues of origin (origin tissues, OT), and **(d)** normal, non-cancerous target tissues (TT) (**Table S1A**). Except for the one quadruple-paired bulk RNA-seq dataset and the paired single-cell RNA-seq dataset, *all non-cancerous OT and TT used in analyzing cohorts were sourced from the GTEx*. To ensure comparability, each set of analyzed gene expression datasets was normalized in a uniform fashion (**Methods**). We then compared the expression of tumor samples to the expression of the corresponding OT and TT. Similarity was quantified by computing the Euclidean distance between the tumor and normal expression profiles of the primary or metastatic tumors to the median expression of their corresponding normal tissue of origin and target tissues – termed their *transcriptomic distance (TD)* (**Methods**). For each tumor sample’s expression profile (either primary tumor or metastasis), we calculated its *TD ratio* by computing the ratio of its TD to the tissue of origin over its TD to the target tissue. Euclidean distance was used as a measure to compute TD, but we also tested the robustness of our approach by using alternative measures like Spearman’s correlation and cosine similarity-based distance to validate our approach (**Methods**). Furthermore, we computed the TD ratios for 50 key cellular pathways from MSigDB’s hallmark gene sets^21^ in each sample, quantifying which pathways are closer to the origin or to the target tissues than expected by chance, as well as the transcriptomic distances between matched primary and metastatic samples for the 10 main cancer hallmarks^22^ (**Methods**). **Figure 1** provides a visual overview of the TD ratio calculation.

**Figure 1:**
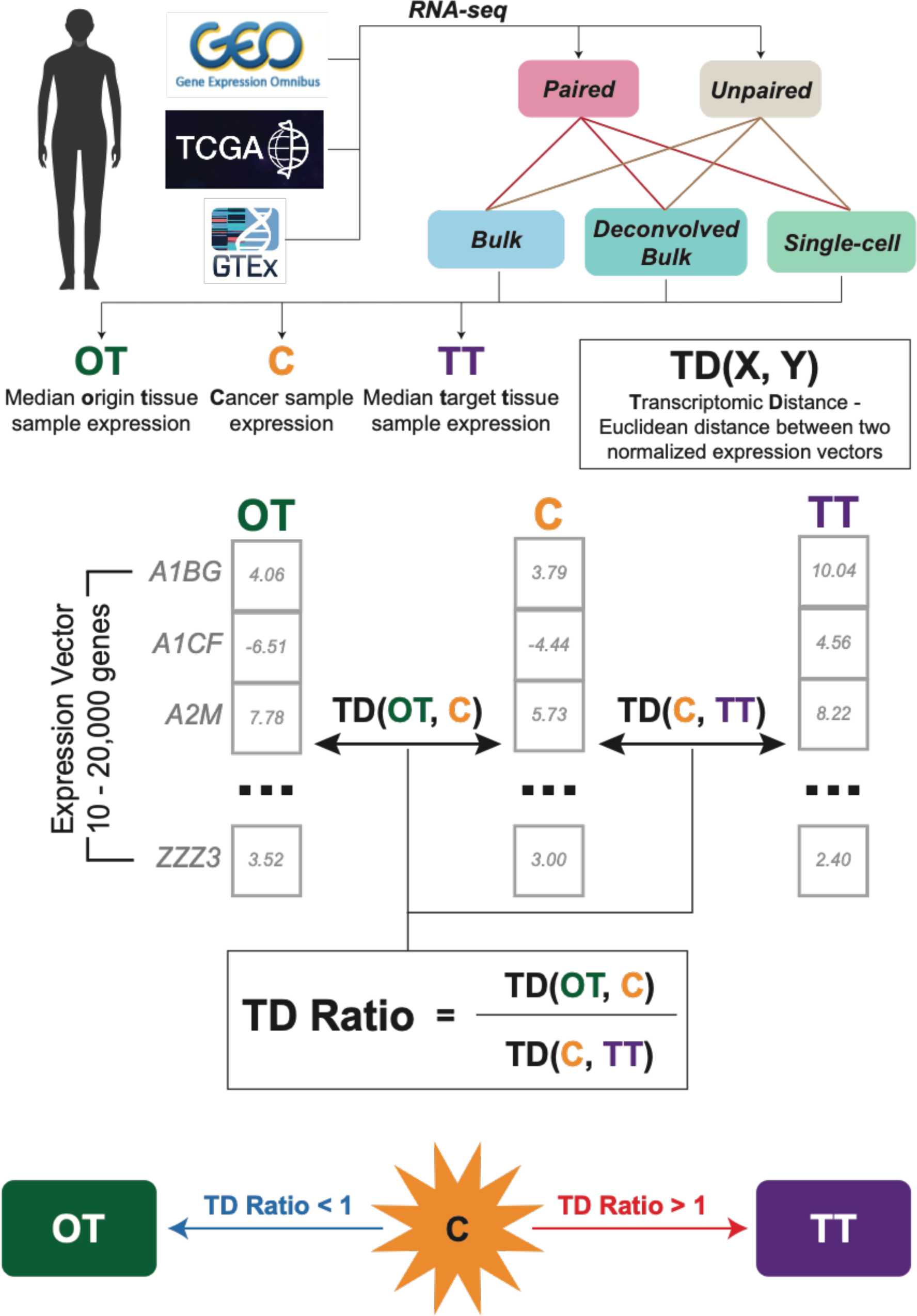
An overview of the approach, depicting the four entities studied in our pipeline: healthy tissue from the primary tumor site, termed origin tissue (OT), primary tumors that have been clinically annotated as metastasizing to a particular target organ, metastatic samples, and healthy target tissue (TT) to which the tumor metastasizes. Data was sourced from several sources and is summarized in **Table S1A**. The transcriptomic distance **TD(X, Y)** of a given tumor sample X denotes the distance of its expression from the median expression of a given origin X and target tissues Y. Finally, the TD ratio denotes whether a cancer sample is more similar to the target tissue (>1), or to the origin tissue (<1). These TD ratios are utilized in the downstream analysis to compare primary and metastatic samples on the whole-genome and pathway levels.

### The landscape of primary and metastatic tumor TD ratios in paired cohorts

To test if paired primary and metastatic tumors are transcriptionally more similar to their origin or target tissue, we begin by examining the TD ratios of primary tumors and their paired metastases from several bulk-paired RNA-seq expression cohorts – all summarized in **Table S1A**. Given the organotropic nature of cancer metastases, we organized our analyses by target tissue, starting with liver metastases.

#### Comparison of paired primary and metastatic tumors expression to origin and target tissues using bulk RNA-seq cohorts

We first analyzed a dataset of 5 quadruple-paired colorectal cancer (COAD) samples, that is, 5 primary colon, 5 metastatic liver, and 5 corresponding OT (colon) and TT (liver) samples (termed *SRR2089755*^14^). While the bulk expression of both primary and metastatic cancers is closer to the OT (TD Ratios < 1), the metastatic cancer expression is significantly closer to the TT than the primary tumor expression (one-sided Wilcoxon Signed Rank test, p = 0.031, **Figure 2A**). This same trend (one-sided Wilcoxon Signed Rank test, p = 0.0059) is found in another paired colon cancer cohort of 9 samples (termed *GSE245351*^23^), where metastases, unlike their paired primaries, exhibited TD ratios closer to and around 1 (**Figure 2A**). These results testify to an expression shift to adapt to the target liver tissue, while the primary cancer’s expression remains more like the origin colon tissue.

**Figure 2:**
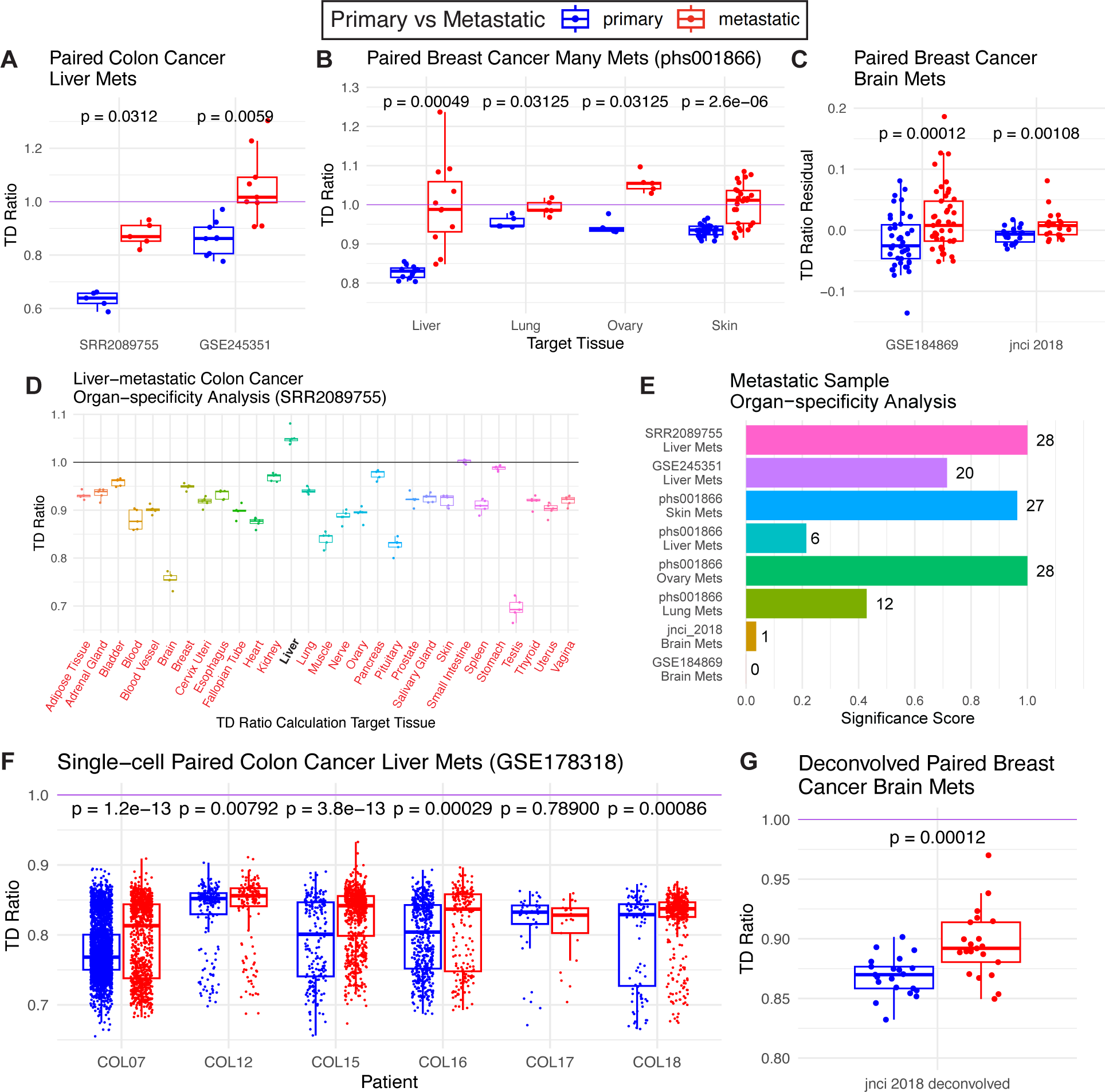
Transcriptomic landscape of primary and metastatic tumors in relation to their corresponding non-cancerous tissues across bulk RNA-seq, paired cohorts. All datasets are summarized in **Table S1A**. Primary sample TD ratios are shown in blue, while metastatic sample TD ratios are shown in red. **(A)** Landscape of TD ratios for the SRR2089755^14^ (n = 5) and GSE245351^23^ (n = 9) COAD liver metastases datasets. **(B)** The same as **(A)** but shows the phs001866^24^ BRCA multiple-site metastases (n = 46). **(C)** The same as **(A)** but shows the GSE184869^25^ (n = 45) and jnci 2018^26^ (n = 21) BRCA brain metastases dataset. **(D)** TD ratios calculated using SRR2089755 samples and various target tissues (TT) sourced from GTEx and notated on the x-axis (**Methods**). The true liver TT is bolded while any other TT marked red has TD ratios significantly lower (one-sided Wilcoxon Signed Rank test, FDR < 0.1) than the true liver TT. (**E**) Bar plot quantifying what is found in **(D)** and expanding the specificity analysis (**Methods**) to all of the bulk paired datasets, showing specificity score and the number of target tissues with lower TD ratios compared to the true target tissue at the end of each bar. **(F)** TD ratios for primary and metastatic epithelial cells from the single-cell colon cancer liver metastatic GSE178318^27^ dataset (n = 6 patients, with 6589 cells total), stratified by the patient from which the samples were collected. **(G)** The same as **(A)** but showing TD ratios for the epithelial cells of the deconvolved jnci 2018 (n = 21) breast cancer brain metastatic dataset. All p-values were generated by the one-sided Wilcoxon Signed Rank test for these paired datasets, testing if metastatic sample TD ratios were greater than primary ones.

An analogous analysis of 46 paired primary and metastatic samples from breast cancer (BRCA) patients (termed *phs001866*) demonstrated similar findings. These BRCA patients had metastases to the liver (n = 22), lung (n = 10), ovary (n = 10), and skin (n = 50) ^24^. Utilizing the bulk expression of normal breast tissue (OT) and normal liver, lung, ovary, and skin tissues (TTs) from the GTEx, TD ratios were calculated for each primary sample and its paired metastasis (**Methods**). Across all four target tissue sites, the metastatic tumors were closer to their TT then OT when compared to the primary tumors (one-sided Wilcoxon Signed Rank test, FDR < 0.1, **Figure 2B**). The ovary metastases exhibited TD ratios all above 1, indicating an overall higher closeness to the ovary TT than the breast OT. The liver, lung, and skin metastases which all exhibited TD ratios around 1, suggesting a ‘midway’ transcriptomic state between the OT and TT.

As our analysis is based on bulk RNA-seq data containing expression from a variety of cell types, we tested if our results are confounded by differences in the tumor’s malignant cell fraction, also known as its “tumor purity”. Reassuringly, we did not find a meaningful correlation between the estimated tumor purity and TD ratio for the datasets analyzed so far (**Methods**, **Table S1D**). However, analyzing two other paired breast cancer brain metastatic cohorts (n = 45, termed *GSE184869*^25^, and n = 21, termed *jnci 2018*^26^) and calculating the samples’ TD ratios, significant correlations were found between the TD ratios and tumor purity scores (p < 0.05, **Table S1D**). Since tumor purity was a possible confounding factor in the analysis of these cohorts, we explicitly regressed out the tumor purity component from the TD ratio via a linear model and acquired the “TD Ratio Residuals”, which represented the TD ratio values free from tumor purity confounding effects (**Methods**). Despite the need for tumor purity regression, both the GSE184869 and jnci 2018 datasets exhibited the same trends as the cohorts reported above, in that the metastatic TD ratio residuals were higher than the primary samples’ TD ratio residuals (p < 0.05, **Figure 2C**). Moreover, we also repeated our analysis using non-Euclidean distance measures (for computing TD ratio) such as Spearman’s correlation-based and cosine-similarity-based measures, which are more immune to any batch effects between datasets, to obtain similar results, thus showing the robustness of our approach (**Figure S1, Supp. Notes 1**).

Given the organotropic nature of cancer metastases, we hypothesized that the metastatic samples may show transcriptomic adaptations specific towards their target tissue (TT) as opposed to some other non-cancerous tissue. As TD ratio computation involves the gene expression of the target tissue, we could compute random control TD ratios for each sample by examining its distance to other tissues in the GTEx data. Starting with the colorectal cancer (COAD) liver metastases dataset (SRR2089755), we performed a “specificity analysis” in which we calculated TD Ratios for each sample fixing the OT to colon and varying the TT (**Methods**). Supporting our hypothesis, we find that the liver metastases’ TD ratios are higher with the actual target normal liver tissue as the TT compared to using any of the other 28 other normal tissues present in the GTEx as the TT, thus suggesting a specific transcriptomic alteration towards their true TTs (one-sided Wilcoxon Signed Rank test, FDR < 0.1, **Figure 2D**). Extending this to all bulk-paired datasets analyzed so far, we defined a score for each dataset indicating for how many target tissues the TD ratios were greater when using the true target tissue versus the other possible target tissues (**Figure 2D, Methods)**. Interestingly, we found mixed results, where liver metastases from the colon along with ovary and skin metastases from the breast have higher specificity in their transcriptomic alterations towards their respective TTs, whereas breast cancer brain, lung, and liver metastases show low specificity (**Figure 2E**).

#### Reinforcing our findings on the paired bulk RNA-seq analysis using a paired single-cell and deconvolved bulk RNA-seq cohorts

Single-cell RNA-seq provides an opportunity to examine the transcriptomic adaptions of primary tumors and their metastases on the cell-type-level, allowing us to perform our analysis in a tumor-purity free manner and home in on the alterations found in just the tumor cells of cancer samples. To this end, we analyzed a paired COAD single-cell cohort with liver metastases (n = 6 patients, ∼6500 cells, termed *GSE178318*^27^). Normal OT (colon) and TT (liver) comparators were taken from Lee at al. (2020) and the Tabula Sapiens database (Tabula Sapiens Consortium, 2022), respectively. A TD ratio was calculated for each cancer epithelial cell in *GSE178318* using 10,000 most variable genes (**Methods, Figure S2**). For five of the six patients, the metastatic cells expressed more similarly to the target tissue than primary cells (FDR < 0.1), suggesting transcriptomic alterations towards the target tissue in the metastatic cells (**Figure 2F**). Reassuringly, when compared to the quadruple-paired bulk COAD liver metastases dataset, both cohorts share a similar trend in TD ratios between primary and metastatic samples, i.e., metastatic samples have greater TD ratios than primary samples (**Figure 2A**).

Recent computational expression deconvolution methods have also allowed for the imputation of cell-type-specific gene expression from a set of bulk tumors. This process provides another way to analyze the transcriptomic alteration found in tumor cells specifically. The paired BRCA brain metastases dataset jnci 2018, analyzed previously in **Figure 2C**, was deconvolved using both a signature for primary BRCA samples and metastatic BRCA brain metastases via the COnfident DEconvolution For All Cell Subsets (CODEFACS)^28^ (**Methods**). After determining the gene expression specific to the cancer epithelial cells in the cohort, a TD ratio was generated for each sample. Reassuringly, this dataset that was once confounded by tumor purity concerns no longer had significant correlations between sample tumor purity and TD ratio after deconvolution (**Table S1D**). As seen in **Figure 2G**, the brain metastases are significantly closer to the brain TT than the primary breast samples, but both primary and metastatic samples are overall closer to the breast OT (FDR < 0.1).

Across all datasets we see a consistent trend of higher TD ratios in metastatic samples than in the paired primaries. Though the specificity of such adaptation appears to vary depending on the cancer type studied, the overall shift towards to the target tissue in metastases compared to primary tumors remains consistent. Reassuringly, across these datasets, dimensional reduction of the gene expression profiles of samples does not result in clusters separated by primary and metastatic classification (for each cancer type), suggesting these results are due to biological difference and not due to batch effects (**Figure S3**).

### The overall landscape of primary and metastatic tumor TD ratios

Given the typically small and limited size of paired datasets, we further explored large-scale, unpaired, pan-cancer bulk RNA-sequencing cohorts to assess transcriptomic alterations across a broader spectrum of cancer types, while acknowledging the inherent limitations of this unpaired data. This unpaired analysis included: **(a)** *primary tumors* of 10 different cancer types sequenced from patient samples *that have been annotated as metastasizing to various target tissues* (collected from the TCGA, n = 306), **(b)** *Metastatic tumors* of 14 different cancer types biopsied from five main target tissues (collected from the MET500 collection, n = 194), and **(c)** *Normal,* non-cancerous tissue samples, composing both the origin and target tissues (GTEx, n = 5,663). To ensure comparability between the three RNA-seq datasets, they were normalized similarly, such that gene vectors were in the same format during TD ratio calculation (**Methods**).

As before, we calculated and examined the TD ratios of primary and metastatic tumors, the results of which are summarized in **Figure 3**. **Figure 3A** shows TD ratios of the primary and metastatic tumors across several cancer types. Consistent with the paired analysis, the transcriptomes of most primary tumors are closer to their corresponding tissue of origin than to that of their eventual metastatic targets, except for primary uterine carcinosarcoma (UCS) and cervical squamous cell carcinoma (CSEC) that have a more equipoise position between their OT and TT (TD Ratio ∼ 1). The transcriptomes of metastatic tumors generally have an equipoise position, with metastases originating from stomach cancer (STAD) representing the closest to origin (median TD ratio = 0.82) and those from pancreatic cancer (PAAD) representing the furthest from origin (median TD ratio = 1.03).

**Figure 3:**
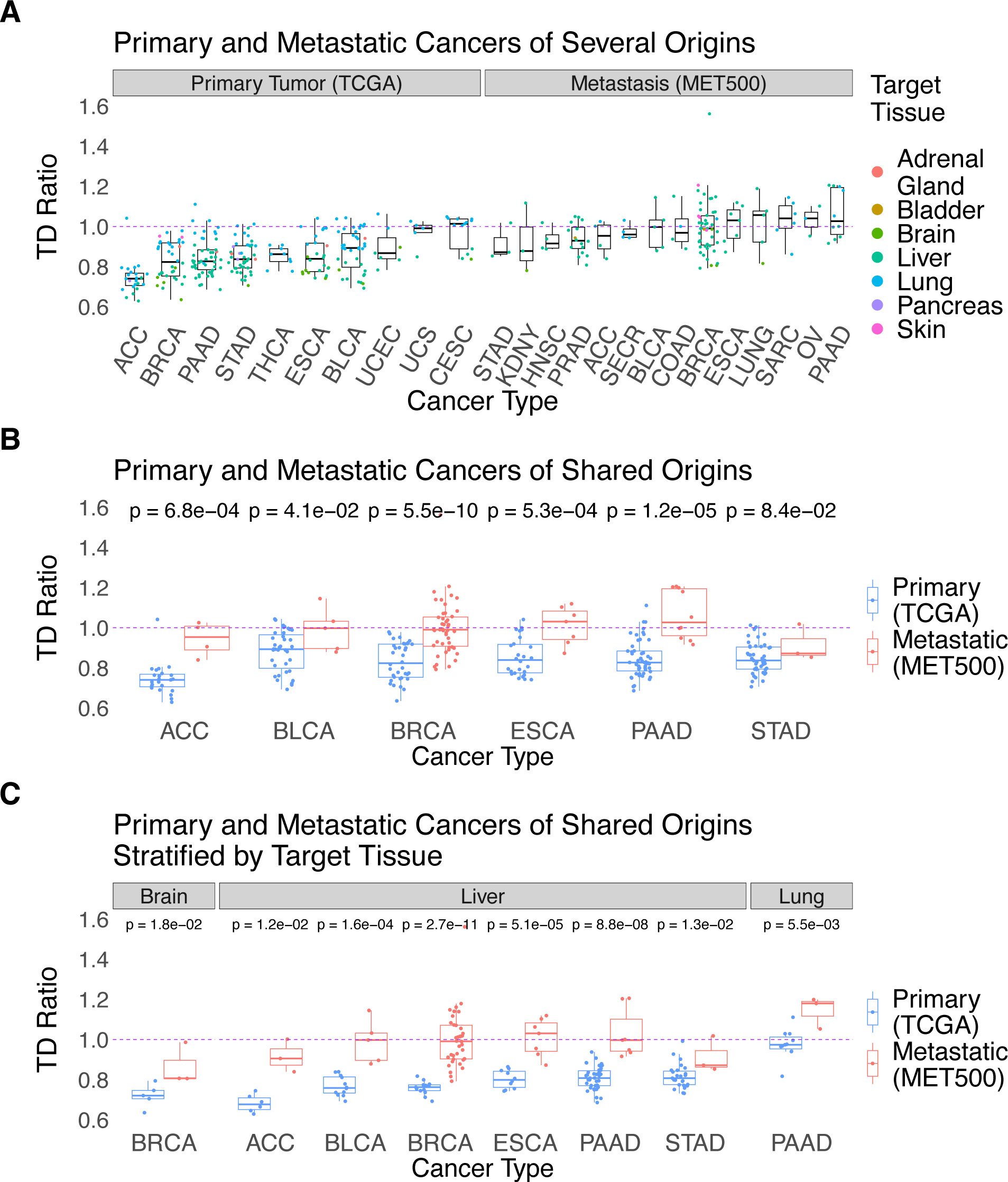
Transcriptomic landscape of primary and metastatic tumors compared to their corresponding non-cancerous tissues across bulk RNA-seq, unpaired cohorts. Tumor samples (primary from The Cancer Genome Atlas^12^, metastatic from MET500^29^) with TD ratios above 1.0 are closer to the target tissue, while those with TD ratios below 1.0 are closer to the origin tissue; the purple line marks the equipoise value of 1.0. In (**A)**, the TD ratios (y-axis) are plotted for all primary and metastatic samples of a given cancer type and ordered by TD Ratio. In (**B)**, all samples are plotted for cancer types with at least 3 samples in both datasets (for any target tissue). Panel **(C)** is the same as **(B)** except the samples are stratified by target tissue. The cancer types in the x-axis are mentioned as TCGA or MET500 abbreviations and their full forms are provided in **Table S1E** (e.g., COAD = colorectal cancer). All p-values presented are the result of one-sided Wilcoxon rank-sum tests for these unpaired datasets. FDR value < 0.1 in all cases presented in **(B)** and **(C)**.

In all six cancer types for which we had sufficient data for primary and metastatic tumors of the same cancer type, the metastatic tumors are closer to the target tissue than the primary tumors that have been reported to have eventually metastasized to that target tissue (FDR < 0.1, **Figure 3B**, **Methods**). This same trend is found when considering only primary samples metastasizing to the brain, liver, or lung (we had most samples for these three target tissues) versus metastases to those sites. The brain and lung metastases subsets only had two primary cancer types with sufficient numbers of samples to make a comparison (breast and pancreatic cancer, respectively), while the liver metastases subset had enough samples from six primary cancer types (**Figure 3C**). In all cases, the metastatic tumors adhere to the aforementioned trend compared to the primary tumors (FDR < 0.1). These results are not confounded by tumor purity (**Methods**, **Table S1D**). Furthermore, the key findings remained similar when replacing the Euclidean distance method for calculating TD ratios with Spearman’s correlation-based and cosine similarity-based distance measures (**Supp. Notes 1**, **Figure S4**).

### Pathways analysis and cancer hallmark activities for different cancer types and target tissues

We next performed a higher-resolution analysis to compute the TD ratios at the pathway level to identify key biological pathways in primary tumors and metastases whose gene expression either shifts towards that of the TT or remains similar to the OT. Using these ratios, we were able to compute the median pathway-specific TD ratios separately across primary and metastatic samples for each cancer type. Empirical p-values (and FDR adjusted values) for each of these TD ratios were computed using random genes with the same size of that pathway. Furthermore, we introduced a comparative metric, the *Δ pathway-specific TD ratio*, to evaluate the extent by which the pathway-specific TD ratio diverges from the TD ratio computed using all genes (**Methods**). A Δ pathway-specific TD > 1 implies that the median pathway-specific TD ratio for all samples is greater than expected in comparison to the all-gene analysis (i.e., the transcriptome of the pathway genes of the tumor is more similar to the TT in comparison to the all-gene analysis). Δ pathway-specific TD < 1 implies the opposite: the transcriptome of the pathway genes of the tumor is more similar to the OT in comparison to the all-gene analysis (**Figure S5**). Of particular interest were pathways of two main types: (a) those whose *Δ pathway-specific TD ratios that flipped from low (<1) to high (>1) (or vice versa) between the primary and metastatic groups*, marking a significant shift in the pathway’s expression during metastasis, and (b) those that have *high Δ pathway-specific TD ratio in primary cancers*, potentially marking ways the primary cancer is preparing for metastasis via expression alteration from its OT.

#### Pathway-specific alterations point to key adaptive transcriptomic changes in metastases

First, we analyzed liver metastases from the paired colorectal (COAD) and breast cancer (BRCA) datasets. The *Coagulation* pathway demonstrates this flipped Δ pathway-specific TD ratio behavior, having a low Δ pathway-specific TD ratio in COAD and BRCA primary samples, but then having a high Δ pathway-specific TD ratio in most of the liver metastases, reflecting expression changes to better match the liver TT in the metastatic state (FDR < 0.1, **Figure 4A, Table S2A**). This same finding is extended in the large-scale MET500 dataset, where both *Coagulation* and *Bile Acid Metabolism* pathways in BRCA liver metastases have high Δ pathway-specific TD ratios (FDR < 0.1, **Figure S6A, Table S2A**). Bile acids, many of which are produced in the liver, are known to promote metastasis and invasion in colon cancer^30^, while the activation of the coagulation cascade is known to play an important role in the metastatic process^31^, supporting the notion that these pathway alterations are important in metastasis. Interestingly, liver metastases from lung cancers (LUNG), esophageal adenocarcinomas (ESCA), and adrenocortical carcinomas (ESCA) in the MET500 show similar results for these two pathways, suggesting their importance in cancerous origins beyond the breast and colon (FDR < 0.1, **Figure S6A, Table S4A**). Of further interest in the breast (dataset phs001866), the *Adipogenesis* and *Fatty Acid Metabolism* pathways show a similar high ratio pattern in subsets of skin, liver, ovary, and lung metastases (FDR < 0.1, **Figure S6B, Table S4B**). Lipid metabolism, a metabolic process that involves these two particular pathways, has indeed been shown to be important in the metastatic process for multiple cancer types^32^. These metastases appear to use these pathways to better adapt to the environments of their target tissues.

**Figure 4:**
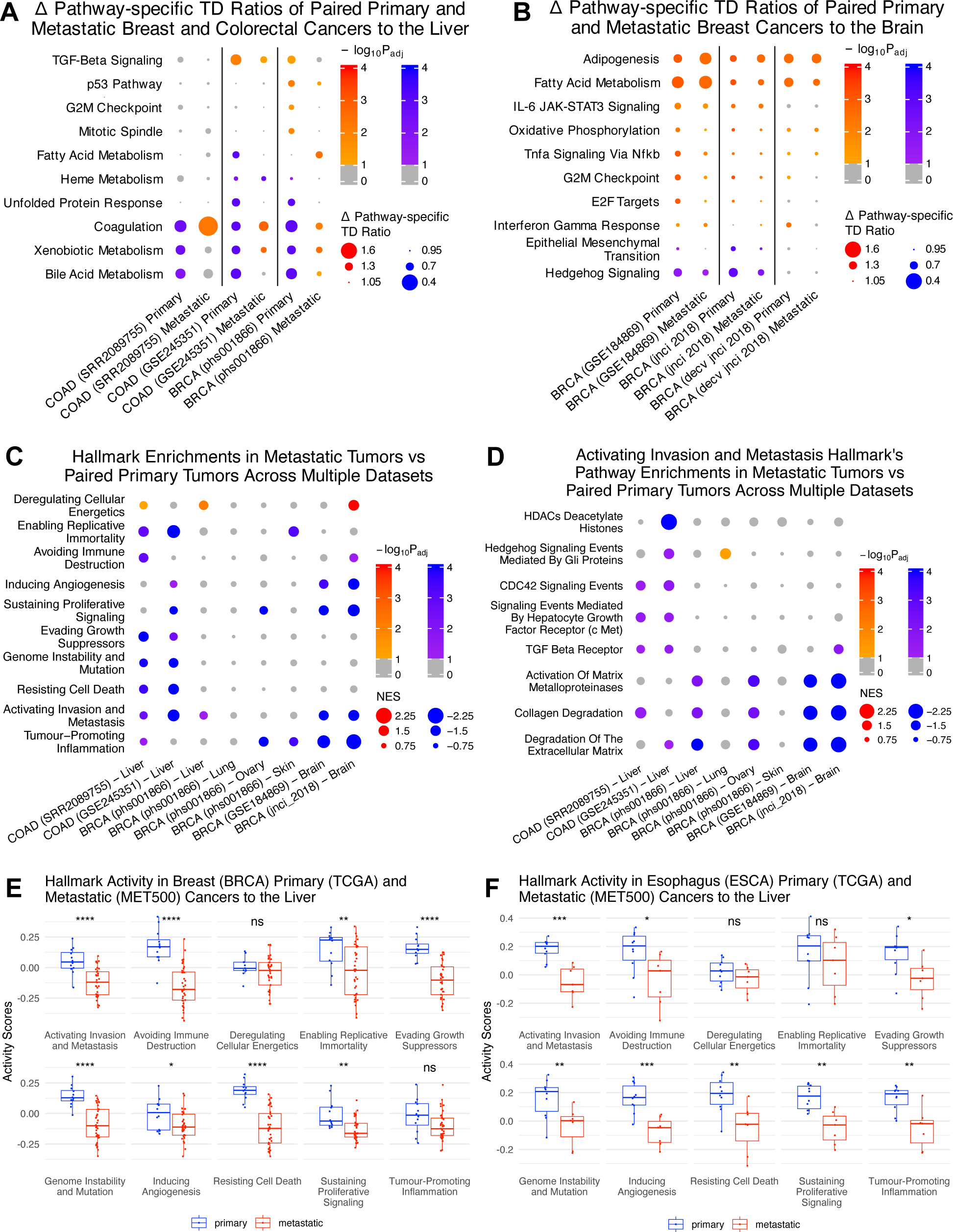
Pathways and cancer hallmarks activity of primary and metastatic tumors. **(A-B)** Heatmap of cohorts with x-axis labelled with dataset and cancer type, and hallmark pathways on the y-axis, where each dot is sized by Δ pathway-specific TD ratio’s distance from the equipoise value of 1. The dots are colored by significance after empirical p-value calculation and false-discovery rate correction (**Methods**). More richly colored dots imply that a TD ratio is further from 1 than expected by chance, i.e., more significant, with insignificant TD ratios (FDR < 0.1) colored in grey. Specifically, **(A)** shows liver metastases from the SRR2089755 (COAD), GSE245351 (COAD), and phs001866 (BRCA) datasets, while **(B)** shows breast cancer brain metastases from GSE184869, jnci 2018, and deconvolved (“decv”) jnci 2018. All Δ pathway-specific TD ratio data pertaining to this pathways analysis can be found in **Tables S2A-B** (one table per subfigure). Not all pathways are shown here for space conservation; pathways were prioritized based on significance across cancer types for plotting (**Methods**). **(C)** Heatmap of pathway enrichments from GSEA run on various datasets (delineated on the x-axis using the format “cancer type (dataset) – target tissue”) and for various pathways shown on the y-axis. The dots are sized by the GSEA normalized enrichment scores (NES), and colored red if the metastatic samples were enriched (NES > 0) or blue if the primary samples were enriched (NES < 0). More richly colored (more red/blue) dot imply a more significant enrichment (FDR < 0.1 as threshold). **(D)** Same as **(C)** but using pathways showing enrichment that compose the “Activating Invasion and Metastasis” hallmark. All NES and FDR values pertaining to **(C)** and **(D)** can be found in **Table S3A** and **Table S3B**, respectively. **(E-F)** GSVA activity scores for the same cancer hallmarks for primary (blue) and metastatic (red) samples breast **(E)** and esophageal **(F)** cancer, respectively.

#### Across cancer types, primary tumors already show early alterations to better match their target tissues of metastasis

We observed that *Fatty acid metabolism* and *Adipogenesis* are not only altered in metastatic samples, but also primary cancers. In breast cancer primaries with both paired and unpaired brain metastases, these two metabolic pathways have high Δ pathway-specific TD ratio, indicating more closeness to the brain target tissue (TT). This suggests that these lipid metabolism pathways are critical in all stages of the BRCA brain metastasis process, that is, before and after metastasis (FDR < 0.1, **Figure 4B, Table S2B, Figure S6B, Table S4B**). Consistent with our findings, lipid metabolism is known to be necessary for proper formation of brain metastases^3^. Along with these lipid metabolism pathways, of interest here is the *oxidative phosphorylation* pathway, which has been strongly correlated with brain metastasis risk in lung cancer patients^33^. *Oxidative phosphorylation* shows greater similarity to the brain TT than the breast OT in our deconvolved breast cancer brain metastases (FDR < 0.1, **Figure 4B**), suggesting the pathway’s critical role in brain-metastatic formation in cancer types beyond just the lung. These primary tumors appear to be adapting to the target tissue before they even reach it.

In addition to the hallmark pathways from MSigDB, we considered the changes between paired primary and metastatic samples for the 10 principal hallmarks (gene sets) of cancer proposed by Hanahan & Weinberg^22^. To determine if each hallmark was more active in primary or metastatic samples, we applied gene set enrichment analysis^34^ (GSEA) on each dataset for gene lists composing those hallmarks^35^ (**Methods**). Colon cancer samples saw primary tumor enrichment in 8 (dataset SRR2089755) and 7 (dataset GSE245351) of 10 hallmarks, while the two breast cancer brain metastatic datasets saw primary tumor enrichment in 4 of 10 shared hallmarks. The breast cancer dataset with four target tissues saw the least significant enrichments, averaging only 2 enriched hallmarks per target tissue (NES < 0, FDR < 0.1, **Figure 4C, Table S3A**). Surprisingly, nearly all of these cancer hallmarks, when significantly enriched, are more active in the primary samples than their paired metastases. Only the *Deregulating Cellular Energetics* hallmark is enriched in the metastases (NES > 0, FDR < 0.1, **Figure 4C, Table S3A**). These hallmarks of cancer delineate critical functions of cancer cells agnostic of cancer type or target tissue, yet we find their overall activities to be lower in metastases, suggesting a decrease in use post-metastasis. When further investigating the *Activating Invasion and Metastasis* hallmark by running the same GSEA on the 28 individual pathways composing it, we found once again that the pathways were mainly enriched in primary samples (**Figure 4D, Table S3B**). This surprising result is not due to some inherent bias in the data as no selective enrichment was found towards primary or metastatic samples when using random gene sets (**Figure S7A, Supp. Notes 2**). Thus, these findings indicate that primary samples, as suggested by their higher levels of *Activating Invasion and Metastasis* activity, are being primed for the metastatic process, after which such processes are less utilized.

Extending this analysis to the large-scale primary (TCGA) and metastatic (MET500) sample cohorts, we performed gene set variation analysis^36^ (GSVA) of the cancer hallmarks on liver metastatic samples and primary samples from cancer types shared between the two cohorts. As with the paired cohorts, the majority of the hallmarks showed higher activity scores in primary tumors, particularly for *Activating Invasion and Metastasis*, than in metastatic tumors of the same cancer type and target tissue (two-sided Wilcoxon rank-sum test FDR < 0.1, **Figures 4E-F**; details provided in **Supp. Notes 3, Figures S7B-E**). Overall, we find that metastatic samples, though known to retain features of their origin tissues^13,19,32^, surprisingly show lower hallmark activities after they leave their tissue of origin.

## DISCUSSION

This analysis is the first to our knowledge to perform a large-scale, pan-cancer transcriptomic landscape analysis comparing primary tumors, metastatic tumors, and normal tissues of tumor origin and target. Although prior studies have attempted such comparisons, they have been limited to one cancer type or do not compare the cancers with both relevant non-cancerous tissues. We first studied the transcriptomic alterations in a small but quadruple-paired cohort of bulk RNA-seq samples, representing the most clinically relevant case. From there, we analyzed several paired primary and metastatic cohorts, and then, given the paucity of paired, multi-cancer datasets available, we used several of the largest publicly available datasets to define aggregate datasets to study expression similarity at the level of cancer type and tissue, integrating bulk, single-cell, and bulk deconvolved data types. Finally, we examined transcriptomic alterations on the pathway and cancer hallmark level, dissecting changes that in both primary and metastatic tumors.

Overall, we find that metastatic tumors across cancer types generally have an equipoise transcriptomic similarity between their tissue of origin and their metastatic target tissue, and significantly greater similarity to the target tissue than the similarity of the corresponding primary tumors of the same type. In contrast, we found that primary tumors, overall, are largely transcriptionally similar to their tissue of origin, which concurs with the current notion that primary tumors remain functionally akin to their tissues of origin in many cancer types^8,19^. The transcriptomic shift in metastases may also be specific to the target tissue of metastasis as shown in the colon cancer liver metastases and breast cancer skin and ovary metastases datasets -- a result in line with the organotropic nature of metastasis. Strikingly, despite the different target tissues and datasets analyzed across the bulk, deconvolved bulk, and single-cell levels, the trend of adapted metastases and more conserved primary tumors remained strong.

When comparing the expression profiles of hallmark pathways in principally liver and brain metastases across paired and unpaired as well as bulk and bulk-deconvolved samples, two main trends emerged. (1) Metastases adapt their metabolism and function to better match that of the target tissue. Subsets of colon and breast liver metastases have *Bile Acid Metabolism* and *Coagulation* pathway expression more akin to the liver than to their origin tissues (also found in lung, esophagus and adrenal gland cancers to some degree in the large-scale dataset), and vice versa with the primary samples, while the same is true for *Adipogenesis* and *Fatty Acid Metabolism* in breast cancer subsets of brain, lung, ovary, and skin metastases. This extreme shift in pathway expression implies a need for those functional pathways in the liver environment, thus marking adaptations to that environment. (2) Primary cancers across several cancer types are primed for metastasis. Both breast cancer primary samples and brain metastases feature several pathways that are closer to the target tissue, e.g., *Adipogenesis*, *Fatty Acid Metabolism,* and *Oxidative Phosphorylation*, indicating that those pathways are important in the metastatic process both before and after its occurrence. Gene set enrichment of the *Activating Invasion and Metastasis* cancer hallmark and its composite pathways demonstrates that these primary tumors, though they have not metastasized, do have pathways active that assist in their potential to invade and metastasize. Furthermore, primary tumors are enriched for several cancer hallmarks when compared to metastatic tumors of the same cancer type. In other words, most cancer hallmarks including *Activating Invasion and Metastasis* are more active in primary samples in comparison to their paired metastases suggesting that these primary tumors are primed for metastasis before it actually happens. Overall, accounting for these three trends and targeting or further investigating some of these pathways may help in the effort to impede metastasis formation.

Our study has a few limitations, some which we have addressed. Tumor purity is a potential confounding factor for the comparative transcriptomic analysis in the bulk as some surrounding normal tissues could be a part of the cancer biopsy. Therefore, we carefully checked and controlled for any meaningful associations between tumor purity estimates and TD ratios for both the unpaired and paired datasets. Furthermore, we reinforced the trends presented via corroboration of TD ratios calculated with single-cell and deconvolved bulk RNA-seq – both free of tumor purity issues. Moreover, our pathway-specific analysis identifies pathways that show alterations over the whole transcriptome-based analysis and therefore this analysis is unaffected by tumor purity concerns. Due to the lack of a large-scale transcriptomic dataset containing paired sets across several cancer types, we carried out a significant fraction of the analysis using aggregate, non-paired datasets. To ensure comparability between datasets, each cohort of RNA-seq datasets was normalized so that the data was compared in the same format and analyzed separately. The robustness of our results was also verified by using alternative distance measures, such as those based on Spearman’s correlation and cosine similarity, that are more immune to batch effects in the TD calculation scheme.

Bearing these limitations in mind, this study asked and characterized whether the transcriptome or particular pathway of a tumor sample is closer to the tissue from which it originated or the tissue to which it metastasizes. Studying this question across cancer and data types as well as various target tissues has enabled us to shed some light on this fundamental question on a large-scale manner, highlighting the importance of pathways and gene sets previously unrealized in certain cancer types. More comprehensive answers to this question await the analysis of large transcriptomic datasets of paired primary tumors and their metastases, both on the bulk and single-cell levels. Nevertheless, this novel systematic analysis of the expression landscape of primary tumors and metastases with respect to their non-cancerous origin and target tissues provides a transcriptome-wide, pan-cancer view of cancer tumors’ adaptations to their metastatic niches.

## METHODS

### Expression Data Collection

RNA-seq gene expression data was collected from the Gene Expression Omnibus, Xena Browser, and other sources collected into **Table S1A**. Most were collected in Fragments Per Kilobase of transcript per Million mapped reads (FPKM) or count form, though some paired datasets were sourced in unique formats (e.g., upper-quartile normalized counts from phs001866) as the counts were unavailable. The gene expression data from the non-cancerous sources was always converted to match the cancerous data to which it was compared, for example GTEx count data was converted to the TMM-CPM format of the GSE184869 before comparing expression vectors.

As a special case, the data for SRR2089755 were sourced from the BARRA:CuRDa browser^37^, which contained formatted counts, instead of the original Lee et al^14^ manuscript, which only contained raw FASTQ files.

### Phenotypic Data Collection

Phenotypic data fields of interest generally included information on whether a tumor is primary or metastatic, biopsy sites, and cancer types for all samples, as tabulated in **Table S1B**.

### Sample Filtering

The datasets were trimmed to just the relevant samples, i.e., those for which cancerous, origin tissue, and target tissue gene expression data existed. The “Samples Used” columns in **Table S1A** reflect the samples considered in the study after filtering. Each dataset had to be filtered somewhat differently as each had differently organized phenotypic data. The informative fields/columns used in filtering of the sources’ phenotype tables are given in **Table S1B**.

### Data Processing

To ensure comparability between non-cancerous and cancerous datasets (**Table S1A**), non-cancerous datasets (downloaded as RNA-seq count data) were converted to the formats of the cancerous data. The counts to upper-quartile-normalized (UQ-normalized) counts conversion was done by dividing each sample’s expression by its 75^th^ percentile value. The counts to trimmed mean of m-values (TMM) – counts per million (CPM) conversion was done with the “edgeR” R package^38^. The counts matrix was fed into the “DGEList” function, after which “calcNormFactors” with parameter “method” = “TMM” and “cpm” functions were applied. In the single-cell data, all 10X counts were converted to CPM (more details in the *Single-cell* section below).

Then, in order to make the comparison between tumor samples and normal tissue simpler, median expression vectors were generated for each tissue type present in the non-cancerous datasets (**Table S1A**). For each tissue type and for each gene, independently for each dataset, the median (*mean* in single-cell datasets because of data sparsity) of the gene across all samples of that tissue type was taken. This generated an expression vector with the median transcriptome expression for each non-cancerous tissue.

For each analysis, the list of genes was limited to those in common between all the datasets involved. These gene intersections and corresponding datasets are summarized in **Table S1C**. All gene names were formatted as gene symbols and converted to gene symbols from ENSG ids when necessary using the “biomaRt” package^39^ in R using the package’s standard vignette.

Rarely in the TCGA, cancer samples were marked as having multiple target tissues, in which case new “pseudo-samples” were created for a sample for each of its target tissue. For example, if a sample was marked as having metastases to the liver, lung, and brain, then the sample would be split into three different samples with the same gene expression as the original sample but each marked with only one target tissue (liver, lung, and brain, respectively).

### Sample-Level Transcriptomic Distances (TDs)

For a given cancer sample, we used a transcriptomic distance (TD) to gauge its distance from a given normal tissue. Transcriptomic distances were calculated as follows: for cancer sample X and normal tissue Y, we treat the expression of the genes in X as a vector. Each vector is log2-transformed (if not already log-transformed from the source) and z-score standardized for normalization and fair comparison. Transcriptomic distance is then the Euclidean distance between these two vectors, which provides a measure of the cancer’s similarity to the normal tissue. This is visualized in **Figure 1**. Alternative measures like Spearman’s correlation and Cosine-similarity based distance was also used for computing transcriptomic distances, for the sake of robustness (**Supp. Notes 1**).

### Transcriptomic Distance Ratios (TD Ratios)

We used the transcriptomic distances from the previous step in order to determine whether a cancer was closer to its tissue of origin or the tissue to which it metastasizes (target tissue). For a particular cancer type, a transcriptomic distance was taken from both the origin tissue and the target tissue. The ratio of the distance to the origin tissue over the distance to the target tissue was then taken. This “TD ratio” is a measure of which tissue the cancer is closer to. TD ratios above 1 indicate that the cancer is closer to the target tissue than the origin tissue, and thus is more similar to the former. TD ratios below 1, on the other hand, indicate that a cancer is closer to the origin tissue than the target tissue (visualized in **Figure 1**).

### Specificity Analysis and Specificity Score

To determine if the transcriptomic alterations in metastatic samples were specific to their target tissues, we charted the landscape of TD ratios using different target tissues. Because the GTEx dataset contained 30 distinct tissues, we calculated TD ratios for samples using all of these tissues as the target tissue (TT) in calculation, sans the origin tissue. In a dataset, the TD ratios generated using the actual TT of metastasis were then compared to the TD ratios generated using the other TTs via one-sided Wilcoxon Signed Rank tests, with false-discovery rate correction following (significance threshold of 0.1). From this, the *specificity score* for that dataset was defined as the number of times the TD ratio distribution using the actual TT was significantly higher than the TD ratio distributions using other TTs.

### Tumor Purity and TD Ratio Residuals

Tumor purity was calculated using the “ESTIMATE” R package^40^. For each cancer sample, the package calculated a score that is a stand-in for purity based on a single-sample Gene Set Enrichment Analysis (ssGSEA). The package was then applied to every bulk dataset to obtain Spearman’s correlations between transcriptomic distance ratios and tumor purity – termed purity correlations (**Table S1D**). Purity correlations were calculated for primary and metastatic samples separately in each dataset, and the resulting p-values from the Spearman’s correlations determined if a dataset needed tumor purity correction. Datasets that needed adjustment had their TD ratios corrected through a linear model; specifically, the residuals derived from the linear regression using the model TD ratio ∼ tumor purity replaced the original TD ratios. These residuals were denoted as “TD Ratio Residuals”.

### Single-Cell Data Processing

The single-cell dataset GSE178318^27^ did not come with phenotypic data, forcing us to infer the tumor and cell types of the tumor cell samples. The 10X raw counts were loaded and processed using the “Seurat” R package (Hao et al., 2021), and sample primary/metastatic classifications were differentiated into primary (by the substring “_CRC”), metastatic (by the substring “_LM”), and blood cells. Quality control (QC) was performed by removing samples with high mitochondrial reads (15% threshold) and only keeping samples with number of genes detected (“nFeature_RNA”) and number of molecules detected (“nCount_RNA”) within three standard deviations of the mean. These QC samples were then clustered using highly-variable genes in principal component analysis (PCA) and t-distributed stochastic neighbor embedding (t-SNE), after which the tumor cells (malignant/epithelial) were extracted by only keeping the two clusters that showed high EPCAM expression. Having the counts of just the epithelial cells (already split into primary and metastatic at this point), these were finally converted into log CPM using the “NormalizeData” function, setting “scale.factor” = “1e6”. This protocol was adapted from the Che et al^27^ publication’s own methods section.

For the non-cancerous datasets, the phenotypic information was provided as cell annotations, so only the quality control step was done with the same parameters as above.

Due to the overall sparsity of the single-cell data, the number of genes used in the TD ratios analysis was limited to the 10,000 most variable genes between the non-cancerous and cancer datasets. TD ratios were generated per patient for different numbers of variable genes, and 10,000 was chosen because that many genes adequately determined TD differences between primary and metastatic samples. More details are provided in the supplementary materials (**Figure S2**).

### Deconvolution Analysis

To extract the epithelial/tumor cell gene expression from bulk data, we needed to first generate a signature of genes delineating each cell type. The dataset deconvolved was jnci 2018, a paired breast cancer (BRCA) primary and brain metastatic dataset, therefore signatures needed to be generated for BRCA primary samples and BRCA brain metastatic samples, independently. Each signature was generated by feeding annotated single-cell data (CPM format) into the CIBERSORTx^41^ online portal, using default parameters, which was then fed in conjunction with the bulk data into a high-performance computing cluster implementation of the COnfident DEconvolution For All Cell Subsets (CODEFACS) ^28^, also using the default parameters. The output gene x sample matrix was filtered to just keep epithelial cells per sample keeping all genes, and it was used for downstream analysis.

For the primary breast cancer data source, we derived a signature matrix of 1,400 genes and nine cell types using a single-cell RNA-seq cohort from Wu et al^42^. These nine cell types included B-cells, cancer associated fibroblasts (CAFs), cancer epithelial cells (malignant), endothelial cells, myeloid cells, normal epithelial cells, plasmablasts, perivascular-like cells (PVL), and T-cells. Meanwhile for the metastatic breast cancer data source, a signature matrix of 3,319 genes and nine cell types using specifically the breast single-cell RNA-seq profiles from Gonzalez et al^17^.These metastatic cell types were malignant cells, vascular smooth muscle cells (vSMCs), mesenchymal stem cells (MSCs), metastasis-associated macrophages (MAMs), endothelial cells (ECs), pericytes (PCs), astrocytes, T cells, and dendritic cells.

### Pathway Selection

Hallmark Pathways were downloaded from MSigDB^21^. These are well-defined pathways reflecting specific metabolic, signaling, or immune states. For each analysis, only pathways with sufficient overlap in the shared genes between the pathway and the dataset being analyzed were kept. In the bulk, a sufficiency threshold of 90% overlap was used, and in the single-cell data due to sparsity a 70% was used instead.

### Pathway-specific TD Ratios

By creating a TD ratio with just the genes in a pathway rather than the entire transcriptome, we can determine if a certain phenotypic feature of a cancer is more similar to the target tissue or the origin tissue. To accomplish this, for a given pathway and tumor sample, we took the genes in that pathway and created expression vectors – one for the tumor sample, and a median expression vector for the origin and target normal tissues of that tumor. Each tumor then had the transcriptomic distance from the origin and target tissues measured using only the genes for just that pathway. Just as before, the tumor sample’s TD ratio was created by dividing the TD to the origin tissue by the TD to the target tissue. The pathway-specific TD ratio was calculated as the median TD ratio of samples of that cancer type. Only combinations of cancer type, primary/metastatic classification, and target tissue with at least three samples were considered here.

### Δ Pathway-specific TD Ratios

These were calculated by dividing the pathway-specific TD ratio (generated for a particular cancer type, primary/metastatic classification, and target tissue, in each pathway) by the median of the TD ratios generated with the same cancer type, primary/metastatic classification, and target tissue, but using all genes (referred to as the *all-genes TD Ratio*). Thus, these Δ pathway-specific TD ratios represented if a pathway was closer to an origin/target tissue compared to the overall transcriptome’s tendency for a given set of samples. This process is summarized and visualized in **Figure S5**.

### Empirical p-value Generation and Correction

Empirical p-values were calculated for each pathway as follows: for 10,000 iterations, a random set of genes the same size as the pathway was selected. The Δ pathway-specific TD ratios were then calculated for these random sets, whereafter the empirical p-value (*p*) was calculated as such: let *r* be the number of times the random set of genes’ ratios were larger than or equal to the ratio for the specific pathway and let *n* be the number of random gene sets (10,000 here).

Then, *p* = (*r* + 1) / (*n* + 1), in accordance with the protocol suggested by North et al^43^. P-values were adjusted by setting them to minimum of *p* and 1 – *p*, as a Δ pathway-specific TD ratio below (more similar to origin tissue) or above (more similar to target tissue) those from the random gene sets could be significant, and FDR (multiple hypothesis)-corrected^44^. In this way, the empirical p-values represented how significant the shift in expression was for any pathway in a cancer type from the origin tissue to the target tissue, or vice versa.

### Acquisition of the Cancer Hallmark Gene Sets

The 10 main cancer hallmarks, originally proposed by Hanahan & Weinberg^22^ in 2011, were mapped to gene sets by Iorio et al^35^ in 2018, consisting of a total of 374 orthogonal pathway gene-sets. The gene sets were all taken directly from that publication and utilized in this work.

### Gene Set Enrichment Analysis on Cancer Hallmarks

Gene set enrichment analysis^34^ (GSEA) was performed on paired datasets to determine if the hallmarks were more enriched/active in primary or metastatic samples. For any given paired dataset, (1) fold changes were calculated for all shared genes between the primary and metastatic (metastatic divided by primary) samples using the unlogged/not log-transformed gene expressions. Then, (2) for each hallmark gene set, to account for the rare instance of distinct genes in the sets and not in the cancer expression, we removed genes not in the cancers’ expression datasets. Finally, after setting the random seed for consistency, (3) we fed the pruned pathways and fold changes into the “fgseaMultiLevel” function from the ‘fgsea’ R package^45^ using 10000 iterations (50000 for rare instances where 10000 resulted in blank output and a suggestion to do more iterations). Default parameters were used for “fgseaMultiLevel” otherwise. The normalized enrichment scores (NESs) and FDR-corrected p-values were taken from the output of the “fgseaMultiLevel” function for downstream analysis.

Sampling of the GSEA results for random gene sets was also performed using the same three steps as the previous paragraph but for 10000 randomly sampled gene-sets from the cancer sample expression matrix genes – the enrichment results of which are shown in **Figure S7A** and described in **Supp. Notes 2.** Furthermore, to run the GSEA on the individual pathways that composed certain hallmarks, those pathways (also gene sets) were simply substituted in for the hallmarks and ran through the same three steps as above.

### Gene-set Variation Analysis on Cancer Hallmarks

To determine the hallmarks’ activities in the unpaired setting, gene-set variation analysis^36^ (GSVA) was used to compare primary and metastatic samples. The unpaired data, consisting of one large normalized and scaled gene expression matrix, was fed along with the hallmark gene sets into the “gsva” function from the “gsva” R package^36^, setting method = “gsva”. The activity score outputs per sample were then compared via two-sided Wilcoxon rank-sum tests. To check if the sampling scores were biased towards higher values in primary or metastatic samples in general, we calculated scores with random gene sets, as done in the gene-set enrichment analysis (**Supp. Notes 2**).

### Statistical Analyses

To compare the distributions of TD ratios in primary and metastatic samples, one-sided Wilcoxon signed-rank tests (for paired data) and one-sided Wilcoxon rank-sum test (for unpaired data) were used^46^. When applicable, multiple hypothesis correction (FDR) using the using Benjamini & Hochberg method was done to correct p-values^44^. Empirical p-value generation for the pathways analysis is described in a previous paragraph. The distributions of cancer hallmark activities as calculated by the gene-set variation analysis were compared via two-sided Wilcoxon rank-sum tests. All statistical analyses were performed in R.

## CODE/DATA AVAILABILITY

The code used for the main results of this work is provided via NIH datashare at https://hpc.nih.gov/~Lab_ruppin/Primary_vs_Met_project_code_for_reviewers.zip for the sake of reproducibility. We used publicly available datasets for this paper, though the access token for the newer GSE245351 is “cnydcwqyndghdah”. All scripts were developed using latest R version 4.3 and Rstudio 2023.06.

## AUTHOR CONTRIBUTIONS

N.U.N. and E.R. conceived the project and supervised the research work. N.S. and C.C-A. carried out most of the analysis with help from N.U.N., S.S. (Stefano Scalera), V.C. The data were analyzed and interpreted by N.U.N., E.R., N.S., C.C-A., P.S.R., S.S. (Sanju Sinha), F.S., K.W., S.M., E.S., A.P-S., G.B. The paper was mainly written by N.S., N.U.N., E.R. with inputs from all authors. All authors have read, commented, and approved the manuscript.

## ACKNOWLEDGEMENTS

This research was supported in part by the Intramural Research Program of the National Institutes of Health (NIH), National Cancer Institute, and the Center for Cancer Research. This work utilized the computational resources of the NIH HPC Biowulf cluster (https://hpc.nih.gov). The results shown here are in whole or part based upon data generated by the TCGA Research Network: https://www.cancer.gov/tcga. We thank Dr. Kuoyuan Cheng and Dr. Yingying Cao for their help on this project, as well as Premvanti Patel for their aesthetic eye in figure design.

## DECLARATION OF GENERATIVE AI AND AI-ASSISTED TECHNOLOGIES IN THE WRITING PROCESS

During the preparation of this work the author(s) used ChatGPT 3.5 in order to improve readability. After using this tool/service, the author(s) reviewed and edited the content as needed and take(s) full responsibility for the content of the publication.

