## Supplementary Notes and Figures for "Charting the transcriptomic landscape of primary and metastatic cancers in relation to their origin and target normal tissues"

**DESCRIPTION OF THE SUPPLEMENTARY TABLES**

Table S1: summarizes various aspects of the datasets used.

A. Summary of the datasets used as well as their contents, origin tissues, target tissues, samples used, download formats, and sources.
B. Where the phenotype (cancer type, metastatic sites, etc.) data for each dataset came from.
C. How many genes were used in the TD ratios calculations for various datasets.
D. Correlations between the tumor purities and TD ratios of samples in each dataset.
E. The cancer type abbreviations mapped to their full names (e.g., OV → ovarian serous cystadenocarcinoma).

Table S2: the delta pathway-specific TD ratio and corresponding FDR results for

A. Figure 4A (Table S6A), which utilizes the liver metastases in SRR2089755, GSE245351, and phs001866 datasets.
B. Figure 4B (Table S6B), which utilizes the brain metastases in GSE184869, jnci 2018, and deconvolved jnci 2018 datasets.

Table S3: contains the GSEA NES scores and corresponding FDR results for

A. Figure 4C (Table S7A), which utilizes several datasets and the 10 cancer hallmarks.
B. Figure 4D (Table S7B), which utilizes several datasets and the 28 pathways within the Activating Metastasis and Invasion cancer hallmark.

Table S4: contains the delta pathway-specific TD ratio and corresponding FDR results for

A. Figure S6A (Table S8A), which utilizes the liver metastases in the MET500 dataset.
B. Figure S6B (Table S8B), which utilizes the primary and metastatic samples in the phs001866 dataset.
C. Figure S6C (Table S8C), which utilizes the brain metastases in the MET500 dataset.
D. Figure S6D (Table S8D), which utilizes the lung metastases in the MET500 dataset.

### SUPPLEMENTARY NOTES

#### *1. Different distance metrics and transcriptomic distance (TD) ratio computation*

In all of these metrics, a lower distance indicates higher similarity between two gene expression vectors.

Euclidean distance - defined as the mathematical Euclidean distance between two gene expression vectors.

$d_{ED} = \sqrt{\sum_{i=1}^{n}(q_i - p_i)^2}$ , where q and p are two gene expression vectors of length n.

Spearman's Correlation-Based Distance - defined by a simple transformation of the Spearman's Correlation between two gene expression vectors.

$d_{CS} = \frac{(1-s)}{2}$, where s is the Spearman's Correlation Rho (**ρ**) between two gene expression vectors.

Cosine Distance - defined by a simple transformation of the Cosine Similarity between two gene expression vectors.
$d_{CD} = \frac{(1-c)}{2}$, where c is the Cosine Similarity between two gene expression vectors.

Then to calculate the TD ratio for a cancer sample, the distance (using any of these three metrics) between the sample's gene expression vector and its origin tissue's gene expression vector was divided by the distance between the sample's gene expression vector and its target tissue's gene expression vector (**Figure 1** demonstrates this visually). Applying all three distance metrics to four of the paired datasets, no differences in the trends were found (**Figure S1**). This finding extended to the large-scale MET500 and TCGA cohorts as well (**Figure S4**), overall suggesting the TD ratio to be a robust metric for determining if samples are closer to their origin or target tissues.

#### *2. Testing for bias in the gene set enrichment analysis.*

To test if the gene set enrichment analysis (GSEA) or variation analyses (GSVA) were biased towards enriching primary or metastatic tumors for the 10 cancer hallmark gene sets, the methods were run on random gene sets (taken from the cancer samples' genomes) 10000 times per dataset using the same cancer samples as in the main analysis. The size of the random gene sets was set to the average of the sizes of the hallmark gene sets (844 genes).

For the GSEA: after running the sampling on a given dataset, the random gene sets' results were divided by their p-values and normalized enrichment score (NES) scores. Results with significant p-values ($p <= 0.05$) were marked as enriched in metastatic if the $NES > 0$, and primary is the $NES < 0$. Otherwise, if the result was not below the 0.05 threshold, it was marked as neither enriched in metastatic nor primary tumors. In sampling GSEA using the paired cancer datasets, no particular bias was found towards enriching primary or metastatic tumors, with the vast majority of random gene sets across datasets showing no significant enrichment in either primary/metastatic tumors (**Figure S7A)**.

For the GSVA: after running the sampling on the large-scale TCGA (primary tumors) and MET500 (metastatic tumors) in one expression matrix, the random gene sets' results were separated by their GSVA normalized activity scores. Results where the activity scores were not different in primary and metastatic tumors, as determined by Wilcoxon two-sided rank-sum test and FDR correction threshold of 0.1, were marked as not being enriched in either primary or metastatic tumors. If there was a significant difference in activity scores for any given random gene set's GSVA result, the median scores for primary and metastatic tumors were compared, with the higher median indicating which class of tumor was enriched. Sampling the TCGA and MET500 tumors in this way, no particular bias was found towards higher activity scores/more enrichment in primary or metastatic tumors. Of the 10000 random gene sets generated, 3351 were enriched in metastatic samples, 3175 were enriched in primary samples, and 3474 were not enriched at all, suggesting no clear bias towards enriching primary or metastatic tumors.

***3. GSVA on the large-scale, TCGA (primary) and MET500 (metastatic) datasets for other cancer types shows similar trends.***

As shown in **Figures S7B-E**, the other cancer types not shown in the main text also showed similar patterns of higher GSVA hallmark activity in the primary cancer samples compared to the metastatic ones. Interestingly, BRCA and Esophagus (ESCA) cancer primary tumors and liver metastases both showed similar levels of activity in the Deregulating Cellular Energetics hallmark, subverting the “primary sample enrichment” trend like in the paired cohorts in the main text. Additionally, other cancer types including adrenocortical carcinoma (ACC), bladder urothelial carcinoma (BLCA), pancreatic adenocarcinoma (PAAD), and stomach adenocarcinoma (STAD) showed the same trend, with primary samples being enriched in 6, 6, 8, and 6 out 10 hallmarks, respectively.

### SUPPLEMENTARY FIGURES

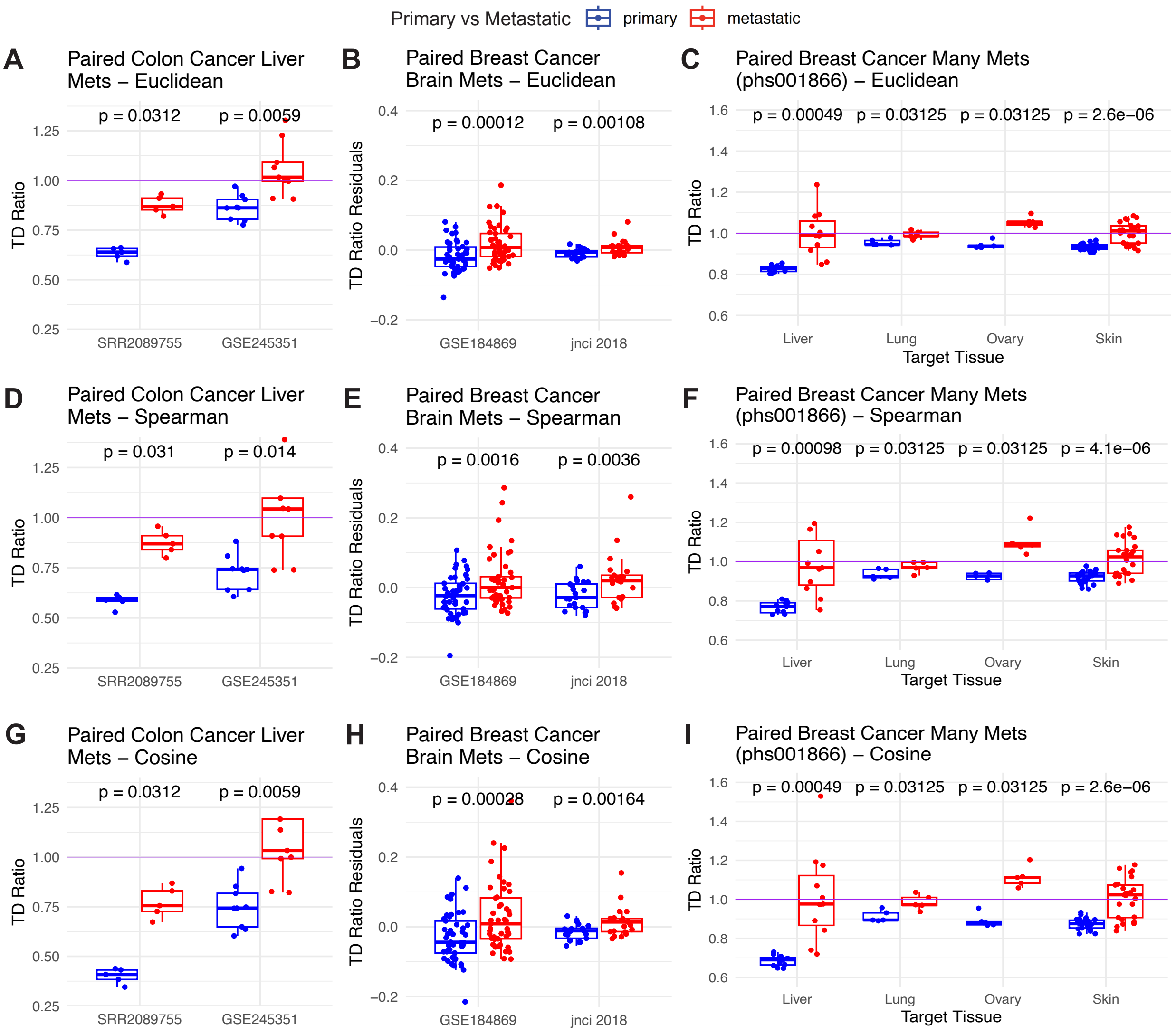


***Figure S1: Comparison of TD ratio distributions when using different distance metrics for paired samples from five different datasets.*** *Three different distance metrics were used to generate TD ratios: Euclidean distance (**A – C**), Spearman's correlation-based distance (**D – F**), and Cosine distance (**G – I**). Each column shows different datasets, with the first showing the SRR2089755 and GSE245351 COAD cancers with liver metastases, the second showing the GSE184689 and jnci 2018 BRCA cancers with brain metastases, and the third showing the phs001866 BRCA dataset with many metastases. All figures use the same color scheme, where primary samples' TD ratios are shown in blue and metastatic samples' TD ratios are shown in red. The datasets being used are shown on the x-axis unless otherwise specified (e.g., with the*

*phs001866 figures whose x axis shows different target tissues). Significance values are from one-sided Wilcoxon signed-rank tests. Overall, using different distance metrics doesn't alter the findings from the TD ratio results, demonstrating the robustness of the ratio. Key word: "mets" = metastases.*

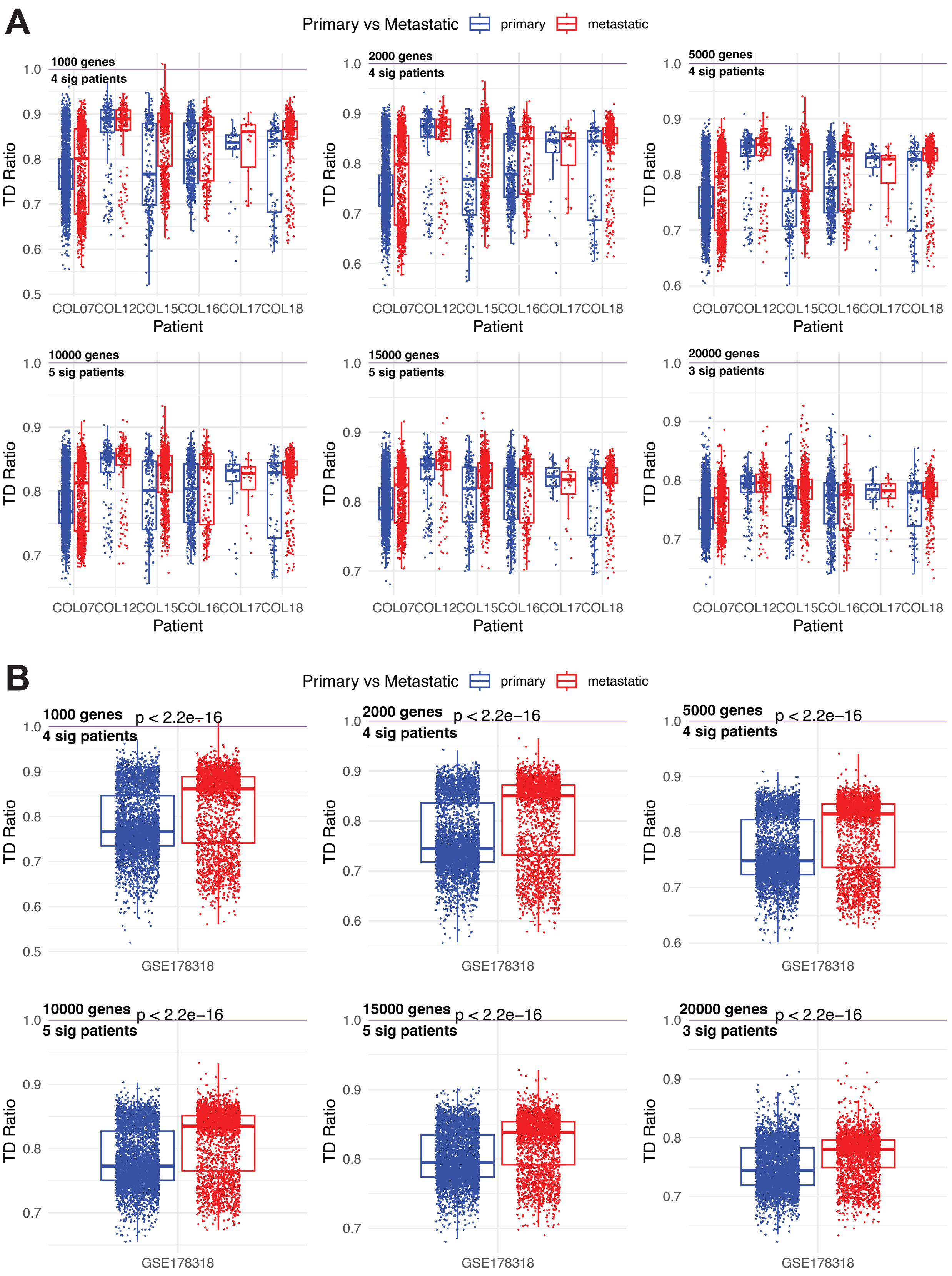
A
Primary vs Metastatic
primary
metastatic
1000 genes
4 sig patients
2000 genes
4 sig patients
5000 genes
4 sig patients
10000 genes
5 sig patients
15000 genes
5 sig patients
20000 genes
3 sig patients
TD Ratio
COL07 COL12 COL15 COL16 COL17 COL18
Patient
B
Primary vs Metastatic
primary
metastatic
1000 genes
4 sig patients
2000 genes
4 sig patients
5000 genes
4 sig patients
10000 genes
5 sig patients
15000 genes
5 sig patients
20000 genes
3 sig patients
p < 2.2e−16
TD Ratio
GSE178318

***Figure S2: TD ratios using different numbers of variable genes in GSE178318 single-cell colon cancer liver metastatic malignant cells. (A)*** *TD ratios for samples grouped by patient and comparisons done between primary and metastatic cells.* ***(B)*** *TD ratios for samples aggregated over all patients. Each plot is annotated with the number of genes used in the TD ratio calculation as well as the number of significant comparisons between primary (blue) and metastatic (red) cells, determined by one-sided Wilcoxon rank-sum tests. Overall, accounting for effect size and significance, the main TD ratios findings are robust to the number of variable genes utilized.*

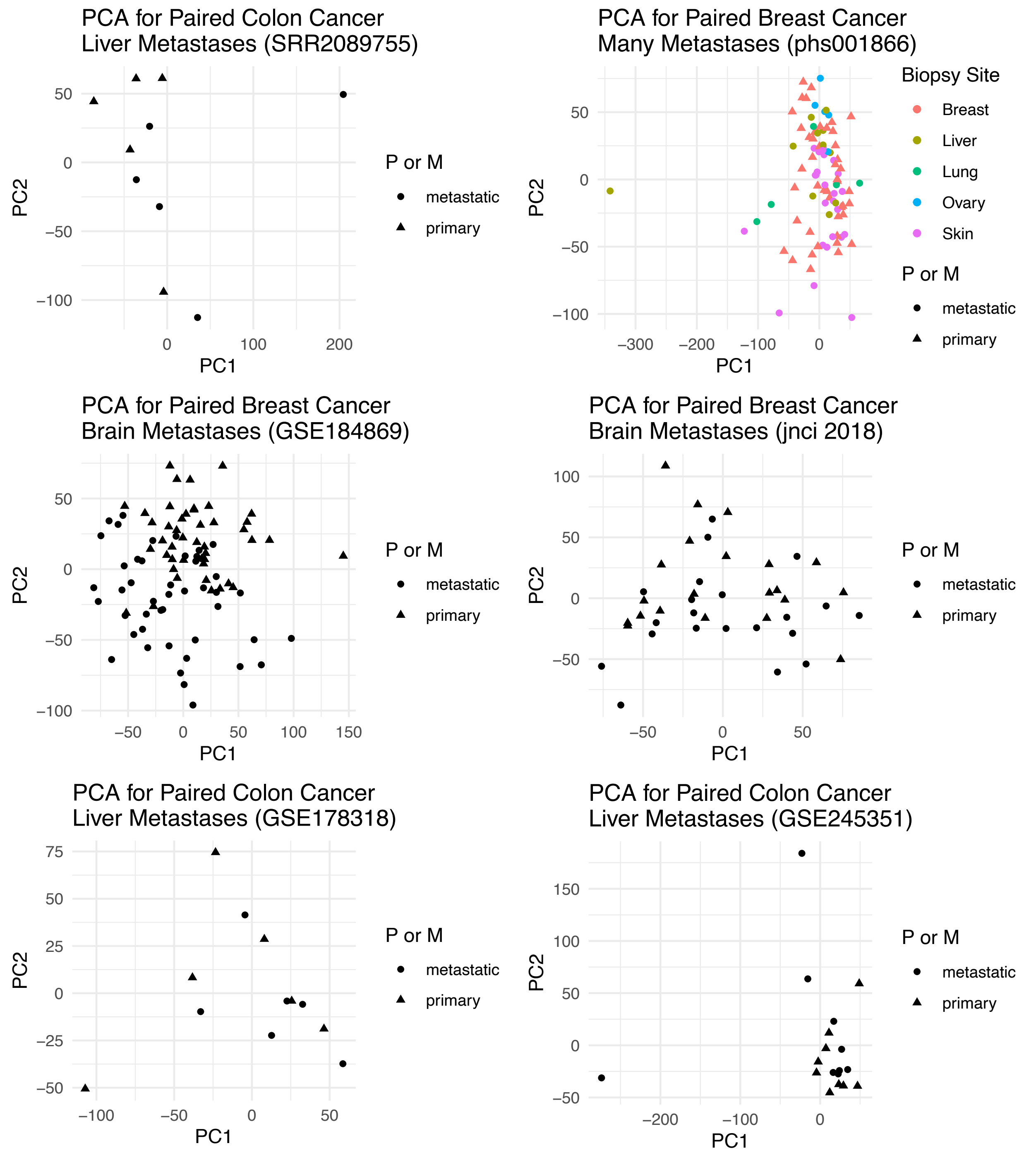


***Figure S3: PCA on datasets.*** *Principal component analysis (PCA) of the gene expression of the paired datasets was performed and the samples were plotted along their first two principal components. As can be seen in all datasets, the samples do not cluster/separate based on primary/metastatic classification.*

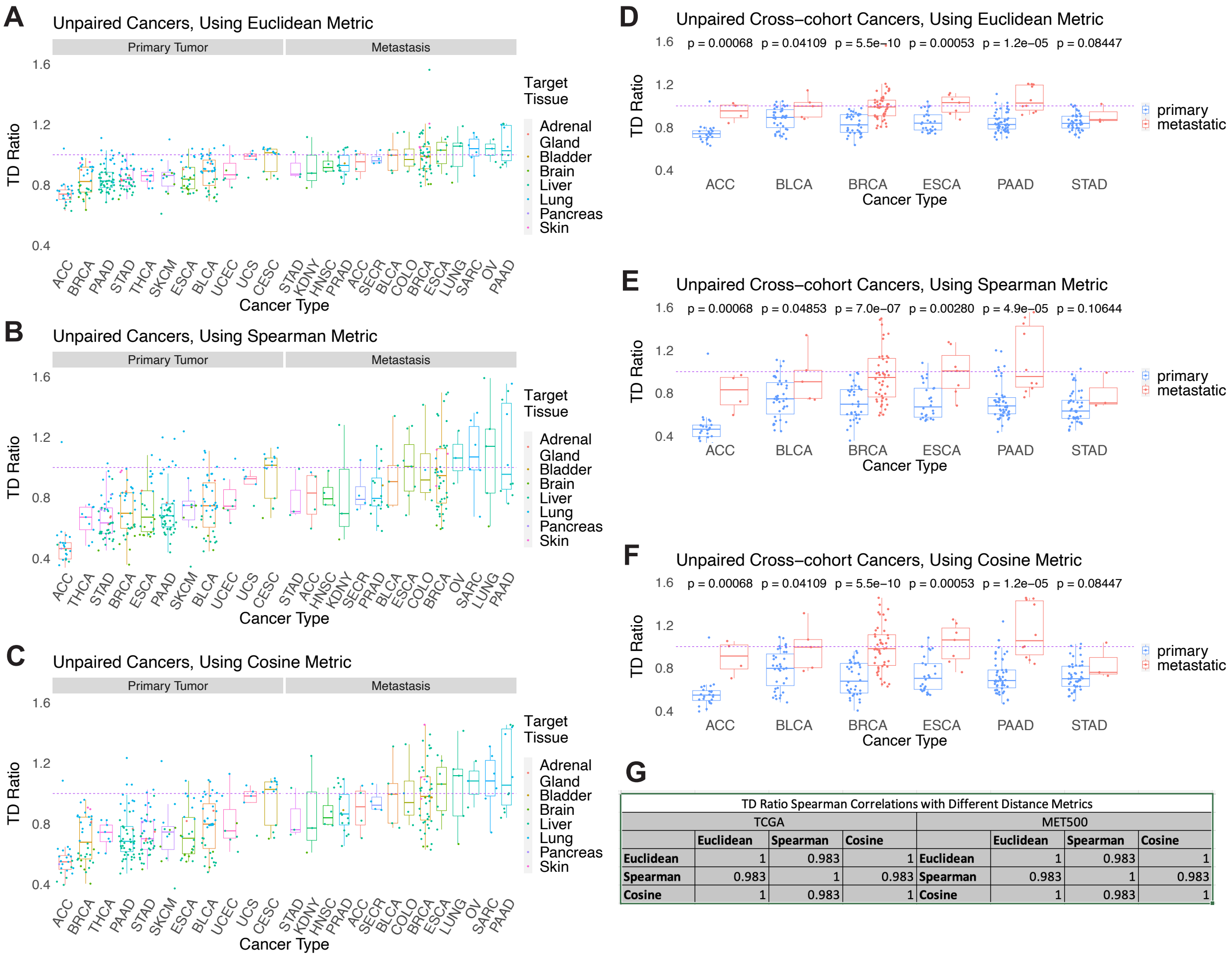


TD Ratio Spearman Correlations with Different Distance Metrics

| | TCGA | | | | MET500 | | |
|---|---|---|---|---|---|---|---|
| | Euclidean | Spearman | Cosine | | Euclidean | Spearman | Cosine |
| **Euclidean** | 1 | 0.983 | 1 | **Euclidean** | 1 | 0.983 | 1 |
| **Spearman** | 0.983 | 1 | 0.983 | **Spearman** | 0.983 | 1 | 0.983 |
| **Cosine** | 1 | 0.983 | 1 | **Cosine** | 1 | 0.983 | 1 |

***Figure S4: Comparison of TD ratio distributions when using different distance metrics in the TCGA and MET500.*** *Three different distance metrics were used to generate TD ratios: Euclidean distance, Spearman's correlation-based distance, and Cosine distance.* ***(A) – (C)*** *The landscape of TD ratios from all samples.* ***(D) – (F)*** *The landscape just using the samples with the same cancer types in both the TCGA and MET500. Distributions are compared via one-sided Wilcoxon rank-sum tests.* ***(G)*** *Spearman's correlations between the TD ratios generated by the different distance metrics, where each entry in the table is Spearman's Correlation Rho. All p-values for the correlations were below 2.2e-16 and were thus not shown. Overall, using different distance metrics doesn't alter the findings from the TD ratio results, further demonstrating the robustness of the ratio. Cancer type abbreviations are provided in* ***Table S1E****.*

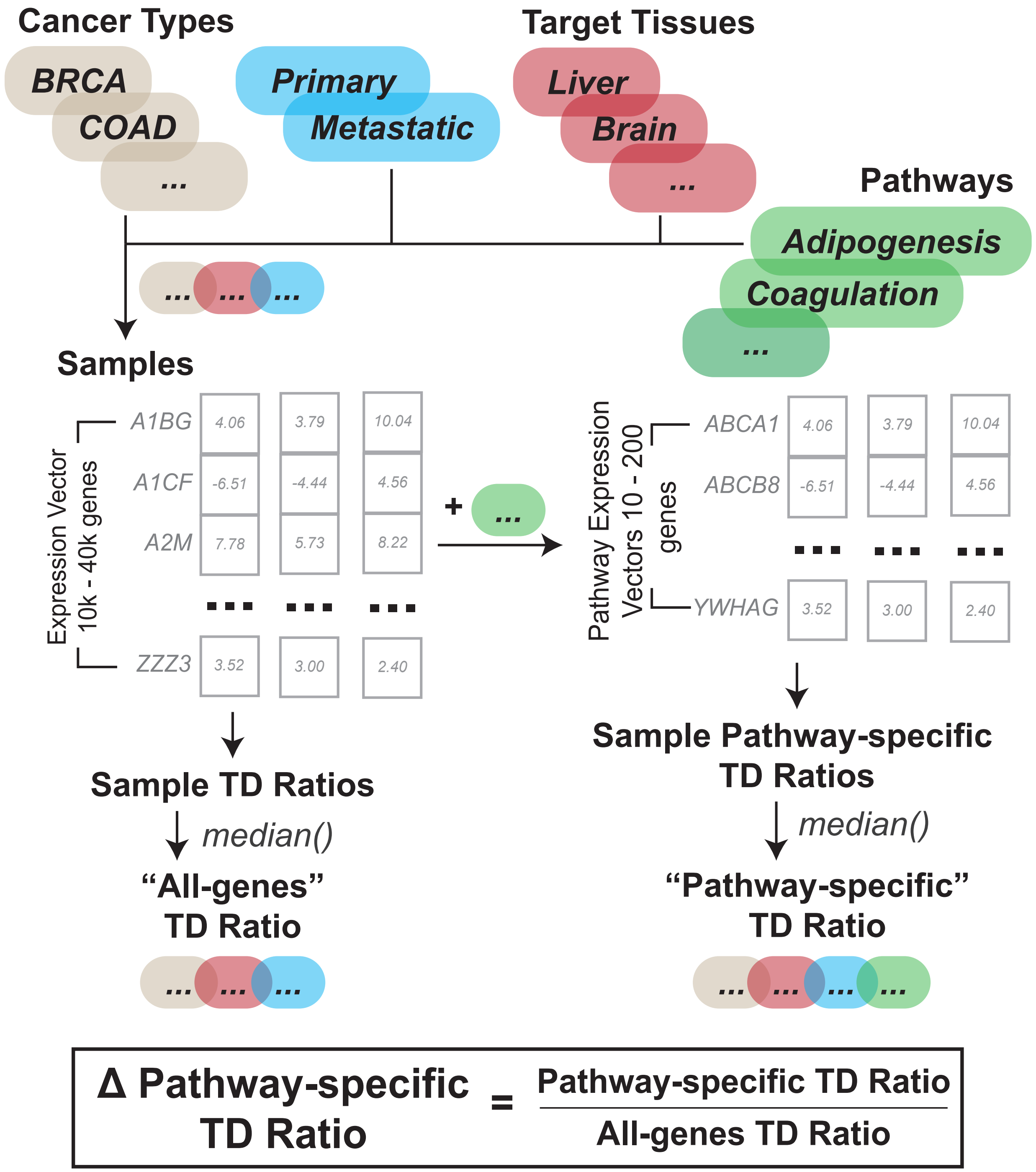


***Figure S5: Calculation of the Δ pathway-specific TD ratio.*** *For a given cancer sample (grouped together by cancer type, primary/metastatic classification, and target tissue) and pathway, two types of intermediate TD ratios are calculated. The "All-genes" TD ratio used the same method as described in* ***Figure 1****, where all of the genes were used in the TD ratio calculations per sample, which were summarized via median into one TD ratio value representing a given cancer type,*

*primary/metastatic classification, and target tissue. Meanwhile, the "Pathway-specific" TD ratio did the same, but only using the genes from the given pathway in the TD ratio calculation. Finally, the Δ pathway-specific TD ratio was calculated as the ratio of the pathway-specific TD ratio and the all-genes TD ratio. Thus, these Δ pathway-specific TD ratios represented if a pathway was closer to an origin/target tissue compared to the overall transcriptome's tendency for a given set of samples.*

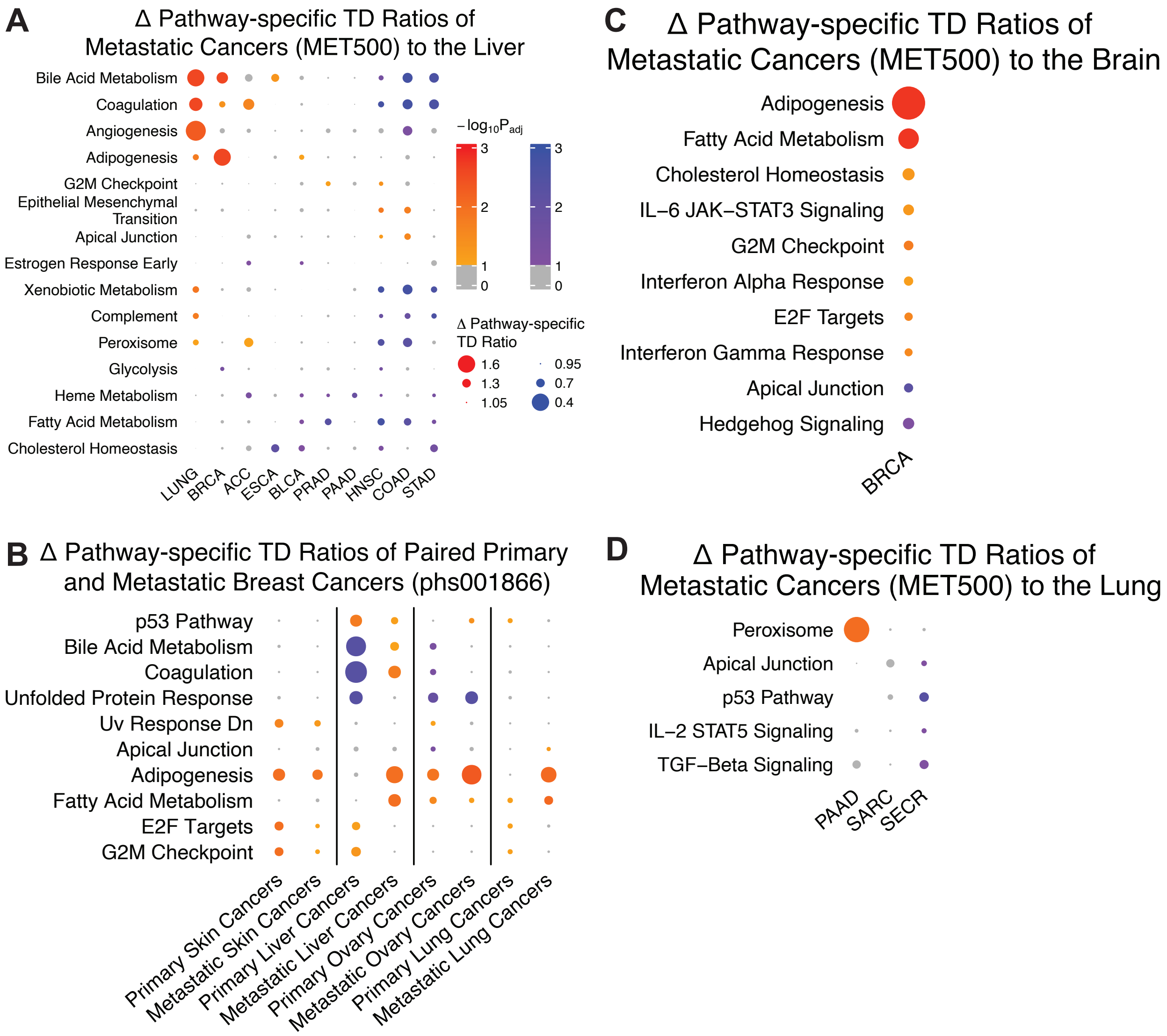


***Figure S6: Pathways analysis of metastatic tumors across multiple cancer types and metastatic sites in the MET500, and all tumors from the phs001866 dataset.*** *Δ pathway-specific TD ratios were generated for metastatic tumors within the MET500 spanning three metastatic sites and*

*several cancer types for which there was sufficient data. We also do the same analysis for all samples within the phs001866 BRCA multiple target tissue cohort (**Methods**). Each subfigure shows a heatmap of cancer types on the x-axis (except for **(B)** which shows the target tissues for various samples within the phs001866 dataset), the association between cancer type label and origin tissue (e.g., COAD = colon adenocarcinoma) is logged in **Table S1E**, and hallmark pathways on the y-axis, where each dot is sized by Δ pathway-specific TD ratio's distance from the equipoise value of 1. The dots are colored by significance after empirical p-value calculation and false-discovery rate correction (Methods). More richly colored dots imply that a TD ratio is further from 1 than expected by chance, i.e., more significant, with insignificant TD ratios (FDR < 0.1) colored in grey. All three subfigures use the legend present in **(A)** and all TD ratio data pertaining to this pathways analysis can be found in **Tables S4A-D** (one per subfigure). Not all pathways are shown here for space conservation; pathways were chosen based on significance for plotting (**Methods**).*

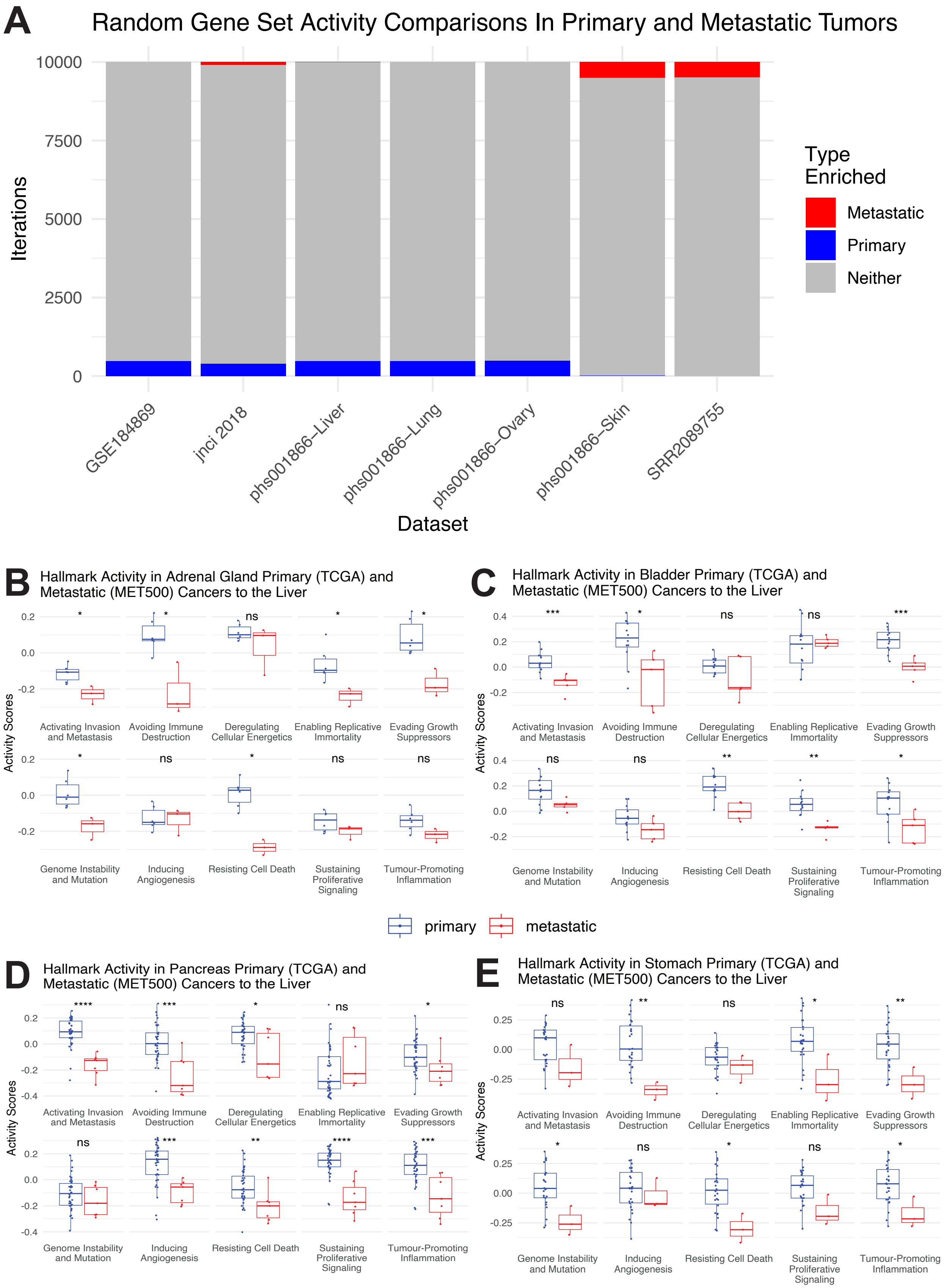
A
Random Gene Set Activity Comparisons In Primary and Metastatic Tumors
Iterations
10000
7500
5000
2500
0
GSE184869
jnci 2018
phs001866–Liver
phs001866–Lung
phs001866–Ovary
phs001866–Skin
SRR2089755
Dataset
Type Enriched
Metastatic
Primary
Neither
B
Hallmark Activity in Adrenal Gland Primary (TCGA) and Metastatic (MET500) Cancers to the Liver
C
Hallmark Activity in Bladder Primary (TCGA) and Metastatic (MET500) Cancers to the Liver
D
Hallmark Activity in Pancreas Primary (TCGA) and Metastatic (MET500) Cancers to the Liver
E
Hallmark Activity in Stomach Primary (TCGA) and Metastatic (MET500) Cancers to the Liver
Activity Scores
Activating Invasion and Metastasis
Avoiding Immune Destruction
Deregulating Cellular Energetics
Enabling Replicative Immortality
Evading Growth Suppressors
Genome Instability and Mutation
Inducing Angiogenesis
Resisting Cell Death
Sustaining Proliferative Signaling
Tumour-Promoting Inflammation
primary
metastatic

***Figure S7: Gene set enrichment analysis bias exploration and gene set variation analysis in various cancer types. (A)*** *Gene set enrichment analysis (GSEA) of random gene sets of size 844 in paired primary and metastatic tumors. GSEA was applied to random sets of genes in the genome of the paired datasets on the x axis, and the enrichment results (metastatic enriched, primary enriched, neither enriched) are counted across 10000 iterations on the y-axis (**Supp. Notes 2**). The samples from the breast cancer dataset with multiple target tissues (phs001866) was split into the four target tissues for analysis. Overall, no bias towards primary or metastatic enrichment is found when using GSEA.* ***(B-E)*** *Gene set variation analysis (GSVA) activity scores for the 10 cancer hallmarks for primary (blue) and metastatic (red) adrenocortical carcinoma, bladder urothelial carcinoma, pancreatic adenocarcinoma, and stomach adenocarcinoma samples, respectively. All p-values are calculated via two-sided Wilcoxon rank-sum tests.*
